# Butyrate producing Clostridiales utilize distinct human milk oligosaccharides correlating to early colonization and prevalence in the human gut

**DOI:** 10.1101/2020.04.15.038927

**Authors:** Michael Jakob Pichler, Chihaya Yamada, Bashar Shuoker, Maria Camila Alvarez-Silva, Aina Gotoh, Maria Louise Leth, Erwin Schoof, Toshihiko Katoh, Mikiyasu Sakanaka, Takane Katayama, Chunsheng Jin, Niclas G. Karlsson, Manimozhiyan Arumugam, Shinya Fushinobu, Maher Abou Hachem

## Abstract

The early life human gut microbiota exerts life-long health effects on the host, but the mechanisms underpinning its assembly remain elusive. Particularly, the early colonization of Clostridiales from the *Roseburia-Eubacterium* group, associated with protection from colorectal cancer, immune- and metabolic disorders is enigmatic. Here we unveil the growth of *Roseburia* and *Eubacterium* members on human milk oligosaccharides (HMOs) using an unprecedented catabolic apparatus. The described HMO pathways and additional glycan utilization loci confer co-growth with *Akkermansia muciniphilia* via cross-feeding and access to mucin *O*-glycans. Strikingly, both, HMO and xylooligosaccharide pathways, were active simultaneously attesting an adaptation to a mixed HMO-solid food diet. Analyses of 4599 *Roseburia* genomes underscored the preponderance of HMO pathways and highlighted different HMO utilization phylotypes. Our revelations provide a possible rationale for the early establishment and resilience of butyrate producing Clostridiales and expand the role of specific HMOs in the assembly of the early life microbiota.

## Introduction

The human gut microbiota (HGM) is a key determinant of health^1–3^. Orthogonal transfer from the mother contributes markedly to the establishment of this community shortly after birth^4,5^. The HGM develops dynamically during infancy until a resilient adult-like community is formed after 2-3 years of life^6–8^. The early life microbiota plays a role in the maturation of the host’s endocrine, metabolic and immune system^9^, and the composition of this consortium appears to be associated with lifelong health effects^10–12^. Therefore, understanding the factors that define the HGM structure during infancy is critical for minimizing the risk for a range of metabolic, inflammatory and neurodegenerative disorders, all associated to specific HGM signatures^13,14^.

Dietary glycans resistant to digestion by human enzymes are a major driver that shapes the developing HGM^6,15^. This is emphasized by the dominance of *Bifidobacterium* in breast-fed infants^7,8^, attributed to the competitiveness of distinct members of this genus in the utilization of human milk oligosaccharides (HMOs)^16,17^. Indeed, the most prominent changes in the infant microbiota occur during weaning and the introduction of solid food^6,7^, whereby bifidobacteria are replaced by Firmictues as the top abundant phylum of the mature HGM. This compositional shift is accompanied by a notable longitudinal increases in the concentrations of the short chain fatty acids (SCFAs) propionate and butyrate (generated from carbohydrate fermentation) during and after weaning^18^.

Butyrate exerts immune-modulatory activities^19^ and is associated with a lowered risk of colon cancer, atherosclerosis, and enteric colitis^20,21^. The bacterial production of butyrate is largely ascribed to Firmicutes *Clostridium* cluster IV and *Clostridium* cluster XIVa that includes members of the *Roseburia*-*Eubacterium* group (Lachnospiraceae family, Clostridiales order), which are abundant and prevalent members of the adult HGM^22,23^. By contrast, the abundance of *Roseburia* spp. is decreased in patients suffering from metabolic, inflammatory and cardiovascular diseases^24–27^. Although butyrate producers are established by the first year of life^27^, the mechanisms underpinning their early appearance (and prevalence) remain unknown.

The evolution of uptake and enzymatic systems that support competitive growth of *Bifidobacterium* species on HMOs^17^ reflects a successful adaptation to the intestines of breast fed infants. We hypothesized that other taxonomic groups, which possess metabolic capabilities that target HMOs, may have an early advantage in the colonization of the infant gut during infancy.

The early emergence of *Roseburia-Eubacterium* members that comprise the main group of butyrate producing bacteria in the human adult gut offers a suitable model group to evaluate this hypothesis. Genomic analyses were suggestive of the presence of putative HMOs utilization gene clusters in *Roseburia* and *Eubacterium* strains. Growth on distinct HMOs (pure or in a complex mixture from mothers milk) and differential proteomics from HMO growth experiments provided compelling evidence for the molecular basis of HMO utilization by this taxonomic group. We corroborated these finding by molecular characterization of the enzymes and transport proteins that confer growth on HMOs. These analyses disclosed an unprecedented enzymatic activity and a previously unknown structural fold highlighting the uniqueness in the enzymology of HMOs utilization by this taxonomic group. We also showed that the unveiled catabolic pathways support cross-feeding on mucin *O*-glycans in co-culture with the model mucin degrader *Akkermansia muciniphila.* Analyses of the metagenome of *Roseburia,* showed a striking conservation and preponderance of the HMO utilization pathways across the genus, underscoring their importance for adaptation to the human gut. This study provides unprecedented mechanistic details into pathways that may contribute to the early colonization and the resilience of key butyrate producing Clostridiales by mediating the catabolism of distinct HMOs and host *O*-glycans.

## Results

### *Roseburia hominis* and *Roseburia inulinivorans* possess conserved gene loci that support growth on distinct human milk oligosaccharides (HMOs)

We hypothesized that HMO utilization may confer an early advantage in the assembly of early life HGM. Genomic analyses of butyrate producers from Lachnospiraceae identified distant homologs of the recently discovered glycoside hydrolase family 136 (GH136) in the Carbohydrate Active enZyme database (www.cazy.org) (Supplementary Fig. 1). This family was assigned based on the lacto-*N*-biosidase LnbX from *Bifidobacterium longum* subsp. *longum* JCM 12 1 7^28^, which cleaves the key HMO lacto-*N*-tetraose (LNT) to lacto-*N*-biose (LNB) and lactose (EC 3.2.1.140; Supplementary Table 1).

We selected two *Roseburia* strains and one from *Eubacterium,* all having GH136-like genes, to examine their HMO utilization capabilities.

Significant growth was observed for *Roseburia hominis* DSM 16839 (*p*<4.0 x 10^-4^) and *Roseburia inulinivorans* DSM 16841 (*p*<1.3 x 10^-4^) after 24 h on media supplemented with HMOs from mother milk, but the growth of *R. inulinivorans* was more efficient (*μ*_max_=0.30 ± 0.01 h^-1^). Next, we carried out growth on building blocks present in HMOs and related oligomers from *O*-glycoconjugates (Fig. 1a-d). *R. hominis* grew efficiently on LNT (*μ*_m_ax=0.22 ± 0.02 h^-1^), its LNB unit (*μ*_m_ax=0.16 ± 0.01 h^-1^) and the mucin derived galacto-*N*-biose (GNB) (*μ*_max_=0.21 ± 0.02 h^-1^). By contrast, *R. inulinivorans* grew better on LNB and GNB relative to LNT. Growth on LNT was also shared by the taxonomically related *Eubacterium ramulus* DSM 15684 from Eubacteriaceae. *R. inulinivorans* was distinguished by growth on sialic acid (Neu5Ac), abundant in HMOs and glycoconjugates (Fig. 1d).

**Fig. 1:**
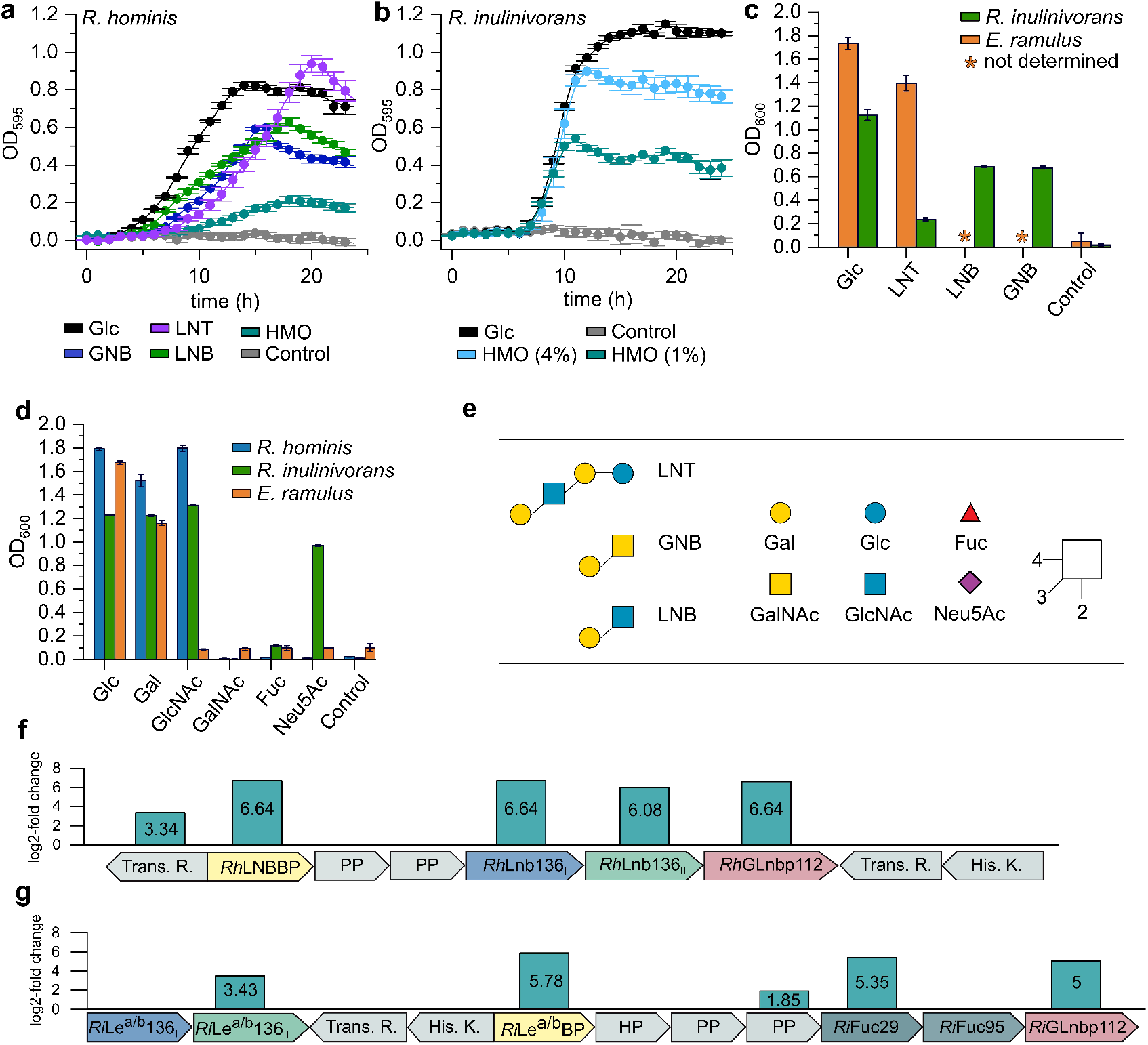
Growth of *R. hominis, R. inulinivorans* and *E. ramulus* on HMOs and upregulation of core HMOs utilization loci: Growth curves of *R. hominis* **(a)** and *R. inulinivorans* **(b)** on glucose, LNT, GNB, LNB, and/or purified HMOs from mothers milk compared to a no-carbon source controls over 24 h. **c**, Growth levels of *R. inulinivorans* on LNT, LNB, GNB and of *E. ramulus* on LNT within 24 h including glucose and a non-carbon source controls. **d**, Growth of *R. hominis, R. inulinivorans* and *E. ramulus* on monosaccharides from HMOs and mucin after 24 h. The growth analyses (**a-d**) on media supplemented with 0.5 % (w/v) carbohydrates (except for *R. inulinivorans* on 1% (w/v) and 4% (w/v) purified HMOs from mothers milk) are means of triplicates with standard deviations. **e**) HMO and mucin-derived oligo- and monosaccharides used for the growth analyses in (**a-d**). The core HMO utilization loci in *R. hominis* (**f**) and *R. inulinivorans* (**g**) identified form proteomic analyses of cells growing on LNT and HMOs from mothers milk, respectively, relative to glucose. The proteomic analyses (**f-g**) were performed in biological triplicates and the log_2_-fold change from the label free quantification of upregulated gene products is shown. Glycan structures presentation according to Symbol Nomenclature for Glycans (SNFG) (https://www.ncbi.nlm.nih.gov/glycans/snfg.html).

To unravel the basis of growth on HMOs, we analyzed the differential proteomes of *R. hominis* and *R. inulinivorans* on LNT and the HMO mixture, respectively, relative to glucose. For *R. hominis* and *R. inulinivorans*, 15 and 62 proteins, respectively, were significantly upregulated (log_2_ fold change > 2). These differential proteomes were dominated by carbohydrate metabolism proteins, especially products of two loci, both encoding an ATP-binding cassette (ABC) transporter, GH112 and GH136 enzymes with putative HMO activities, as well as sensory and transcriptional regulators (Fig. 1f,g). The HMO locus of *R. inulinivorans* is extended with two fucosidases of GH29 and GH95. The specificity-determining solute binding proteins (SBPs) of the ABC transporters of *R. hominis* (*Rh*LNBBP) and *R. inulinivorans* (*Ri*Le^a/b^BP) were the first and fifth top-upregulated proteins in the HMO proteomes, respectively. In addition, the GH112 LNB/GNB phosphorylases were within the top 3 and 12 upregulated proteins in *R. hominis* and *R. inulinivorans*, respectively. In *R. inulinivorans* two additional loci encoding sialic acid and fucose catabolism proteins, were also upregulated (Supplementary Fig. 3).

### Transport proteins of *R. hominis* and *R. inulinivorans* capture HMO blocks and host-derived oligosaccharides

The proteomic analyses highlighted the putative protein apparatus required for growth on HMOs. The solute binding proteins (SBPs) of two ABC transporters in *R. hominis* and *R. inulinivorans* were within the top 8% upregulated proteins, hinting their involvement in uptake of HMOs. Both SBPs recognized distinct HMOs and ligands from host-glycans (Table 1, Supplementary Tables 2 and 3, Supplementary Fig. 4). The *R. hominis* SBP (LNB-binding protein, *Rh*LNBBP) displays a preference for LNB followed by about 3.5 fold lower affinities towards GNB and LNT. By contrast, the SBP of *R. inulinivorans* (Le^a/b^ binding protein, *Ri*Le^a/b^BP) prefers fucosyl-decorated Lewis b (Le^b^) tetraose and Lewis a (Le^a^) triose followed by LNB and GNB, whereas no binding was detected to LNT (Table 1). The loss of the fucosyl unit at the terminal reducing GlcNAc reduced the affinity of *Ri*Le^a/b^BP about 5-fold for blood group H antigen triose type I (H triose type I) relative to Le^b^ tetraose. *Ri*Le^a/b^BP had no affinity for lacto-*N*-neotetroase (LN*n*T) and blood group A antigen triose (A triose). Lactose and 2’-fucosyllactose (2’-FL) were not recognized by either SBP. These results established the capture of specific HMOs and related ligands by these SBPs and the differentiation of their specificities, e.g. preference of *Ri*Le^a/b^BP to fucosylated ligands at the terminal reducing GlcNAc.

**Table 1:**
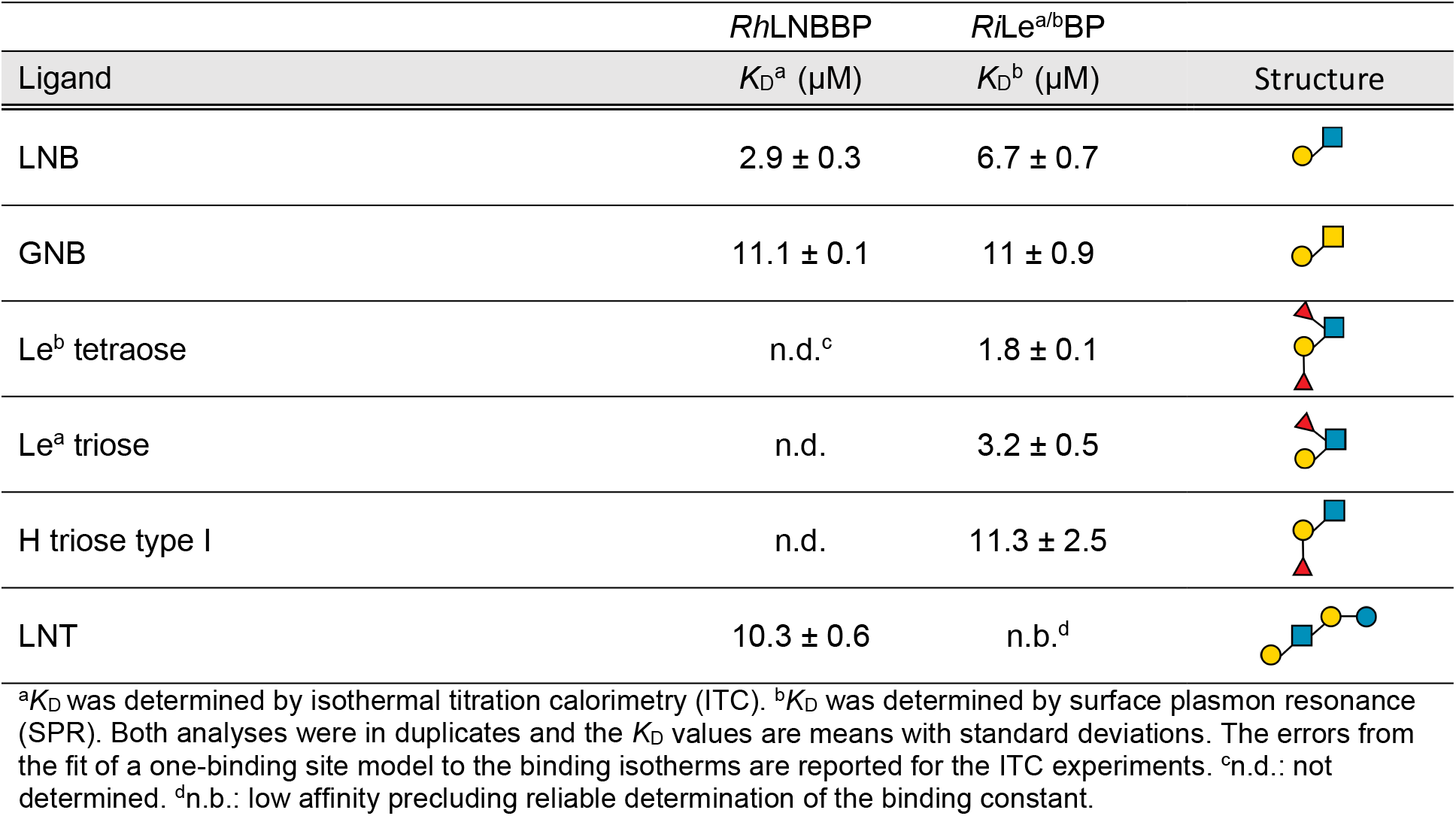
Binding analysis of HMOs and related host-derived oligosaccharides to transport proteins from *R. hominis* and *R. inulinivorans.*

| Ligand | *Rh*LNBBP<br>$K_{\text{D}}^{\text{a}}$ ($\mu\text{M}$) | *Ri*$\text{Le}^{\text{a/b}}$BP<br>$K_{\text{D}}^{\text{b}}$ ($\mu\text{M}$) | Structure |
| --- | --- | --- | --- |
| LNB | $2.9 \pm 0.3$ | $6.7 \pm 0.7$ | |
| GNB | $11.1 \pm 0.1$ | $11 \pm 0.9$ | |
| $\text{Le}^{\text{b}}$ tetraose | n.d.<sup>c</sup> | $1.8 \pm 0.1$ | |
| $\text{Le}^{\text{a}}$ triose | n.d. | $3.2 \pm 0.5$ | |
| H triose type I | n.d. | $11.3 \pm 2.5$ | |
| LNT | $10.3 \pm 0.6$ | n.b.<sup>d</sup> | |

<sup>a</sup> $K_{\text{D}}$ was determined by isothermal titration calorimetry (ITC). <sup>b</sup> $K_{\text{D}}$ was determined by surface plasmon resonance (SPR). Both analyses were in duplicates and the $K_{\text{D}}$ values are means with standard deviations. The errors from the fit of a one-binding site model to the binding isotherms are reported for the ITC experiments. <sup>c</sup>n.d.: not determined. <sup>d</sup>n.b.: low affinity precluding reliable determination of the binding constant.

### Functionally diverse GH136 enzymes confer the initial hydrolysis of HMO blocks and related oligosaccharides within *Roseburia* and *Eubacterium*

The difference in transport preferences between *R. hominis* and *R. inulinivorans* was indicative of different routes for the utilization of HMO-blocks and related oligomers.

The affinity of *Rh*LNBBP from *R. hominis* for LNT, suggested the uptake and subsequent intracellular degradation. Besides the ABC importer, the HMO utilization locus in *R. hominis* encodes distant homologs of the two proteins reported to be necessary for the heterologous expression of the GH136 lacto-*N*-biosidase from *B. longum^29^.* The homologs *Rh*Lnb136_I_ (LnbY in *B. longum)* and *Rh*Lnb136_II_ (Fig. 1f and Supplementary Fig. 1) that harbors the catalytic residues (LnbX in *B. longum)* were highly co-upregulated in the LNT proteome of *R. hominis.* Both proteins lacked predicted signal peptides and transmembrane domains (Supplementary Fig. 8a), in contrast to the *B. longum* counterparts, suggesting the intracellular degradation of LNT in *R. hominis.* Only co-expression and co-purification of *Rh*Lnb136_I_ and *Rh*Lnb136_II_ resulted in an active lacto-*N*-biosidase (henceforth *Rh*Lnb136) (Fig. 2b, Supplementary Table 4). These findings and the observed co-upregulation, suggested that a hetero-oligomer of the *Rh*Lnb136_I_ and *Rh*Lnb136_II_ subunits assembles the catalytically active *Rh*Lnb136. Next, we demonstrated phosphorolysis of LNB and GNB to α-D-galactose-1-phosphate and the corresponding *N*-acetylhexosamines GlcNAc and GalNAc, respectively (Supplementary Fig. 6f), by the GH112 GNB/LNB phosphorylase (*Rh*GLnbp112) in the same locus (Fig. 1f and Supplementary Fig. 1). This enzyme has comparable specific activities for LNB and GNB (Supplementary Table 5) consistent with the growth data on these disaccharides. The functional lacto-*N*-biosidase and GNB/LNB phosphorylase further support the HMO catabolism role of the locus.

**Fig. 2:**
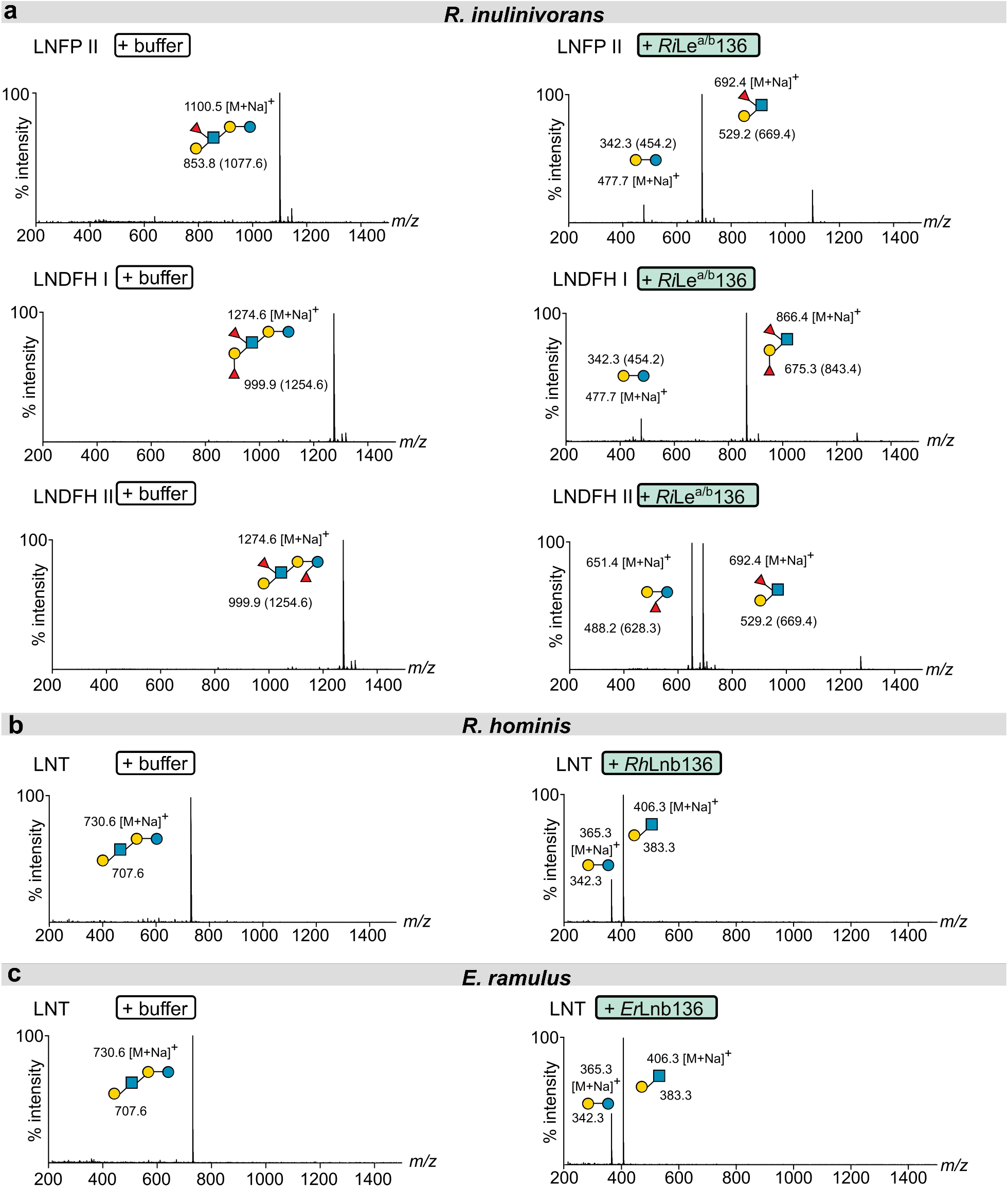
Different specificities of GH136 enzymes in *Roseburia* and *Eubacterium* mediate the degradation of distinct HMOs. **(a)**, Activity of *Ri*Le^a/b^136 on fucosylated HMOs. **(b)**, Activity of *Rh*Lnb136 on LNT. **(c)**, Activity of *Er*Lnb136 on LNT. **(a-c)**, The hydrolysates were analyzed by MALDI-ToF MS without **(b,c)** or with **(a)** previous permethylation. **(a),** Masses of methylated sugars are in parentheses and the ion peaks correspond to the sodium adducts of the methylated sugars. **(a-c)** relative intensity (% intensity) is shown.

The SBP of *R. inulinivorans* had no measurable affinity for LNT in accord with the poor growth (Fig. 1c). Intriguing differences were also observed in the GH136 homolog from the HMO-upregulated locus in *R. inulinivorans* compared to *Rh*Lnb136: 1) the *Ri*GH136_I_ subunit has an N-terminal transmembrane domain, 2) a signal peptide was predicted at the N-terminus of *Ri*GH136_II_, 3) the presence of two C-terminal putative carbohydrate binding modules in *Ri*GH136_II_ (Supplementary Fig. 8a). Co-expression of *Ri*GH136_I_ and *Ri*GH136_II_, lacking the transmembrane domain and signal peptide respectively, resulted in an active enzyme with an unprecedented specificity. This enzyme (*Ri*Le^a/b^136) released Lewis a triose or Lewis b tetraose from fucosylated HMOs including lacto-*N*-fucopentaose II (LNFP II), lacto-*N*-difucohexaose I (LNDFH I) and lacto-*N*-difucohexaose II (LNDFH II) (Fig. 2a and Supplementary Fig. 6a). To our knowledge, enzymatic activity on the glycosidic bond at the reducing end of a fucosylated-GlcNAc unit in the above HMOs has not been reported to date. The products of *Ri*Le^a/b^136 are the preferred ligands for *Ri*Le^a/b^BP, suggesting the uptake of these products by the ABC-transporter. Next, we showed that the concerted action of *Ri*Fuc29 and *Ri*Fuc95 that act on α-(1→4) and α-(1→2)-linked L-fucosyl, respectively mediates the complete defucosylation of Le^b^ tetraose, Le^a^ triose and H triose type I (Supplementary Fig. 6b-d). Initial defucosylation by *Ri*Fuc29 is required for releasing the 1→2 linked L-fucosyl in Le^b^ tetraose by *Ri*Fuc95. Finally, we showed that the GH112 from *R. inulinivorans* (*Ri*GLnbp112) phosphorolyzes LNB and GNB equally efficiently (Supplementary Fig. 6e, Supplementary Table 5).

### A domain with a previously undescribed fold is required for the activity of GH136 enzymes on HMOs

The identification of a *Rh*Lnb136_II_ homolog with high (>60 %) amino acid sequence identity in *E. ramulus* (*Er*Lnb136; Supplementary Fig. 1 and 8a,b) is consistent with the proficient growth of this strain on LNT. Surprisingly, no adjacent GH136_I_ homolog was identified. Instead, the functionally unassigned N-terminal region of *Er*Lnb136 displayed ≈40% amino acid sequence identity to *Rh*Lnb136_I_ (Supplementary Fig. 8b), which was suggestive of the fusion of *Er*Lnb136_I_ and *Er*Lnb136_II_ into a single protein. Indeed, *Er*Lnb136 displayed ≈3.9 fold higher catalytic efficiency on LNT than *Rh*Lnb136 (Supplementary Table 4). A single thermal transition was observed for the unfolding of *Er*Lnb136, suggesting a cooperative unfolding and intimate interaction of the two domains (Supplementary Fig. 8c). We crystallized this enzyme to discern the interactions between the two subunits/domains compulsory for activity within GH136. The crystal structures of selenomethionine (SeMet)-labelled and native *Er*Lnb136 were determined at 1.4 Å and 2.0 Å resolution, respectively (Supplementary Table 6). The C-terminal catalytic GH136 domain (*Er*Lnb136_II_, from AA 242-663) assumes a β helix fold (Fig. 3) similar to the bifidobacterial homolog *Bl*LnbX (Supplementary Table 7). The LNB molecule bound in the active site is recognized by ten potential hydrogen bonds and aromatic stacking of the Gal unit onto W548 (Fig. 3f and Supplementary Fig. 9a). Interestingly, the GlcNAc sugar ring of LNB in *Er*Lnb136 adopts an ^4^*E* conformation (*φ* = 232° and *ψ* = 68°), enabling the O1-OH to adopt a pseudo-axial position to form a direct hydrogen bond with the catalytic acid/base residue (D568) (Supplementary Fig. 9a). Moreover, the D575 O^δ2^ of the nucleophile is positioned appropriately for a nucleophilic attack on the anomeric carbon of the GlcNAc at a distance of 3.2 Å (Fig. 3f).

**Fig. 3:**
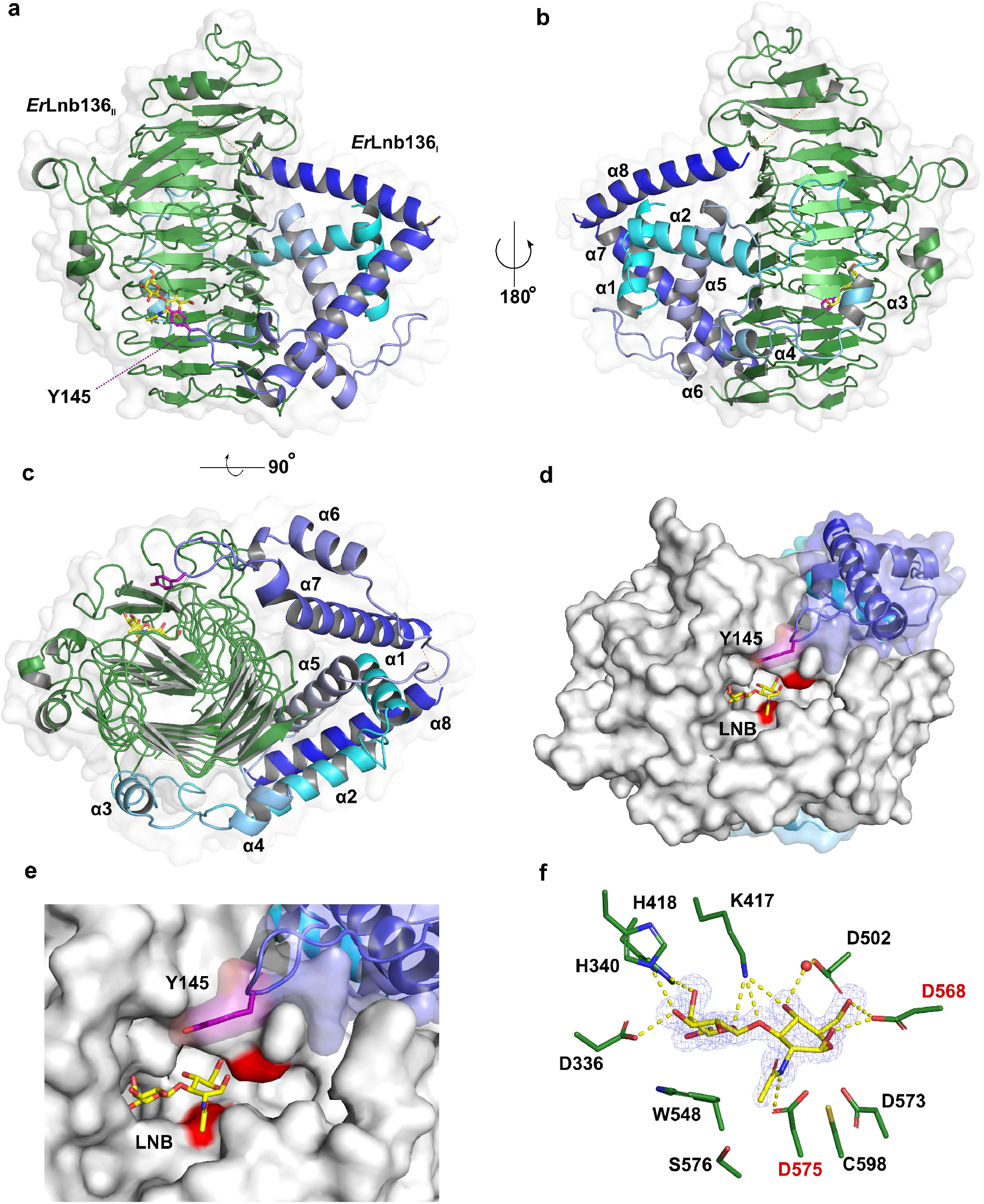
Crystal structure of the GH136 lacto-*N*-biosidase from *E. ramulus* (*Er*Lnb136). **(a-c)** Overall structure and a semitransparent surface of *Er*Lnb136 consisting of an N-terminal domain designated as *Er*Lnb136_I_ (cyan-blue) and a C-terminal β-helix domain (green) -*Er*Lnb136_II_. The enzyme is shown in **(a)** a view orthogonal to the C-terminal β helix domain**, (b)** the view of (a) rotated 180° and **(c)** a view along the axis of C-terminal β helix domain, to highlight the interaction of the *Er*Lnb136_I_ and *Er*Lnb136_II_ domains. **(d)**, A molecular surface top view of the active site and a close up view **(e)** to illustrate the contribution of the N-terminal *Er*Lnb136_I_ domain to the active site architecture, especially the tyrosine (Y145, magenta) that contributes to substrate affinity. **(f)** The weighted *mF_o_-DF_c_* omit electron density map (contoured at 4.0 σ) of the LNB unit (yellow sticks) bound at the active site of *Er*Lnb136 is shown. The water (red sphere) mediated and direct hydrogen bonds that recognize the LNB are shown as yellow dashed lines. **(d-f)** The catalytic nucleophile (D575) and catalytic acid/base residue (D568) are highlighted with red labels. **(a-c)** Disordered regions (residues 180-199 and 225-241) are shown as orange dotted lines

The N-terminal domain (*Er*Lnb136_I_, from AA 7-224) consists of 8 α-helices (α1-α8) (Fig. 3a-c) and assumes a previously unknown fold, stabilized by the central helix α1. The structurally most related protein to *Er*Lnb136_I_, a peptidyl-prolyl cis-trans isomerase with a chaperone activity from *Helicobacter pylori* (5EZ1), shares weak structural similarity restricted to helices α6 and α7 (Supplementary Fig. 9b, Supplementary Table 7). The *Er*Lnb136_I_ domain embraces the sides and back of the β helix domain (Fig. 3a-c). These extensive inter-domain interactions (solvent inaccessible interface ≈1618 Å^2^), stabilize the protein structure with *Δ*G= −17 kcal mol^-1^. Remarkably, the α6-α7 loop of *Er*Lnb136_I_ forms a part of the active site with the solvent accessible sidechain of Y145 positioned near the active site (5.7 Å to the GlcNAc O1 atom of LNB) (Fig. 3d, e). The Y145A mutant showed a 4.9-fold higher *K_M_* (Supplementary Table 4, Supplementary Fig. 8d), suggesting that this residue contributes to substrate interactions, possibly at the +1 subsite.

### Cross-feeding and increased butyrate production from *Roseburia* in mucin cocultures with *Akkermansia muciniphila*

HMOs and *O*-glycans from glycolipids and glyco-proteins including mucin share structural motifs. The high affinity of the SBPs from *Roseburia* for GNB suggested possible foraging of mucin (and/or oligomers from glycoconjugates) and thereby a metabolic interplay of *Roseburia* with mucolytic HGM members. To evaluate possible mechanisms of cross-feeding we compared *Roseburia* growth on mucin with and without the model mucin degrader *A. muciniphila^30^.*

A co-culture of *R. hominis* and *R. inulinivorans* displayed no growth within 24 h on a mucin mixture and only poor growth after 48 h (Supplementary Fig. 7a,b), in contrast to *A. muciniphila* that grew well within 24 h. The co-culture of the two *Roseburia* species and *A. muciniphila* grew to a significantly higher *OD*_600_ than *A. muciniphila* alone (*p* < 3.7 x 10^-6^ at 24 h, *p*< 1.3 x 10^-3^ at 48 h)(Supplementary Fig. 7a). This growth is supported by a 4.5 fold higher butyrate level in the coculture supernatants than *Roseburia* alone (24 h). After 48 h, a slight increase in butyrate concentration was also detected in cultures containing only *Roseburia* consistent with the growth data (Supplementary Fig. 7c).

To unveil the basis for the *Roseburia* growth, the proteomes of *R. hominis* and *R. inulinivorans* were compared between co-cultures of *Roseburia* and *A. muciniphila* grown on mucin and glucose, respectively. For *R. hominis* and *R. inulinivorans,* 31 and 93 proteins, including several CAZymes, were significantly upregulated (log_2_ fold change > 2) (Supplementary Fig. 7e-h) relative to the glucose co-cultures. The transport protein *Rh*LNBBP and *Rh*GLnbp112 from the *R. hominis* HMO locus (Fig, 1f) were the top 6^th^ and 10^th^ most upregulated proteins in the mucin proteome of *R. hominis,* respectively, highlighting the role of this locus in cross-feeding on host glycans (Supplementary Fig. 7g). In *R. inulinivorans,* the corresponding proteins *Ri*Le^a/b^BP and *Ri*LNBBP were also significantly upregulated with log_2_ fold changes of 2.77 and 4.74, respectively. However, the top upregulated protein in the *R. inulinivorans* proteome was a SBP of an ABC transporter colocalised with genes encoding a blood group A- and B-cleaving endo-β-(1→4)-galactosidase (*Ri*GH98), a putative α-galactosidase of GH36 and an α-L-fucosidase (GH29), which was the top fourth upregulated protein in the mucin proteome of *R. inulinivorans* (Supplementary Fig. 7h). The upregulation of this locus suggested that *R. inulinivorans* possesses a functional machinery for directly accessing certain mucin oligomers. We expressed the predicted extracellular *Ri*GH98 and demonstrated robust release of blood group A and B oligomers from mucin and related *O*-glycans (Supplementary Fig. 7i-j, Supplementary Table 9). The co-upregulation of a locus encoding a fucose utilization pathway (Supplementary Fig. 3a) is in accordance with the release of fucosylated oligomers by *Ri*GH98. Another route of foraging, was suggested by the high upregulation of the sialic acid catabolism pathway (Supplementary Fig. 3b), which likely confers the potent growth of *R. inulinivorans* on this substrate (Fig. 1d). These findings establish that the HMOs utilization machinery and additional functional operons support co-growth with *A. muciniphila* on mucin.

### The HMO utilization loci are preponderant in the *Roseburia* genome

The HMO loci, defined by the co-occurrence of GH136 and GH112 genes, are conserved in 5 *Roseburia* reference genomes (Supplementary Fig. 1). To broadly examine the structure and conservation of these loci, the presence of homologs of the aforementioned genes was mapped across 4599 previously reconstructed *Roseburia* genomes^31^. As a reference signature for a central catabolic pathway, the presence of GH10 xylanase genes, compulsory for xylan utilization as shown in *R. intestinalis*^32^, was also analyzed. Strikingly, the GH112 and GH136 HMO utilization genes are about 2-3 fold more abundant than the GH10 counterparts (Fig. 4a), indicative of the broader distribution of the HMO loci compared to the xylanase locus, which is mainly conserved in *R. intestinalis*. The GH136_I_ and GH136_II_ genes have a similar abundance, which is about 30 % lower than that of GH112. This overall trend is reiterated from analyses of individual species-level genome bins (SGBs), but differences in the co-occurrence patterns of GH136 and GH112 genes in different *Roseburia* phylotypes were observed (Fig. 4b). Interestingly, GH136 genes were either absent or far less abundant than GH112 counterparts in several SGB, whereas the converse was only true for a single SGB (4959) that represents 0.65% of the total number of the analysed *Roseburia* genomes. The higher abundance of GH112 genes, associated with the phosphorolysis of HMO or mucin derived disaccharides, prompted us to analyze the organization of 1397 loci, defined by the presence of GH112 sequences in the same metagenomic dataset with a more stringent threshold (70% identity to GH112 sequences present in 5 *Roseburia* reference genomes (Supplementary Fig. 1)). The composition of the gene landscapes appeared to be SGBs specific (Fig. 4c). An ABC transporter, GH136_I_/GH136_II_, and transcriptional regulators were, however, the most frequently co-occurring genes with GH112 gene, which offers a robust signature of the core HMO utilization loci (Fig. 4d) and validates the broad distribution of the pathways described in the present study. Additional CAZymes and carbohydrate metabolic genes were also frequently co-occuring in the vicinity of GH112 genes, suggesting that additional glycan utilization capabilities are clustered around the HMO loci. The *Ri*GH136-like sequences (Fig. 4d) are likely to be underestimated due to the divergence of this clade of GH136 that resembles a previously unknown specificity.

**Fig. 4:**
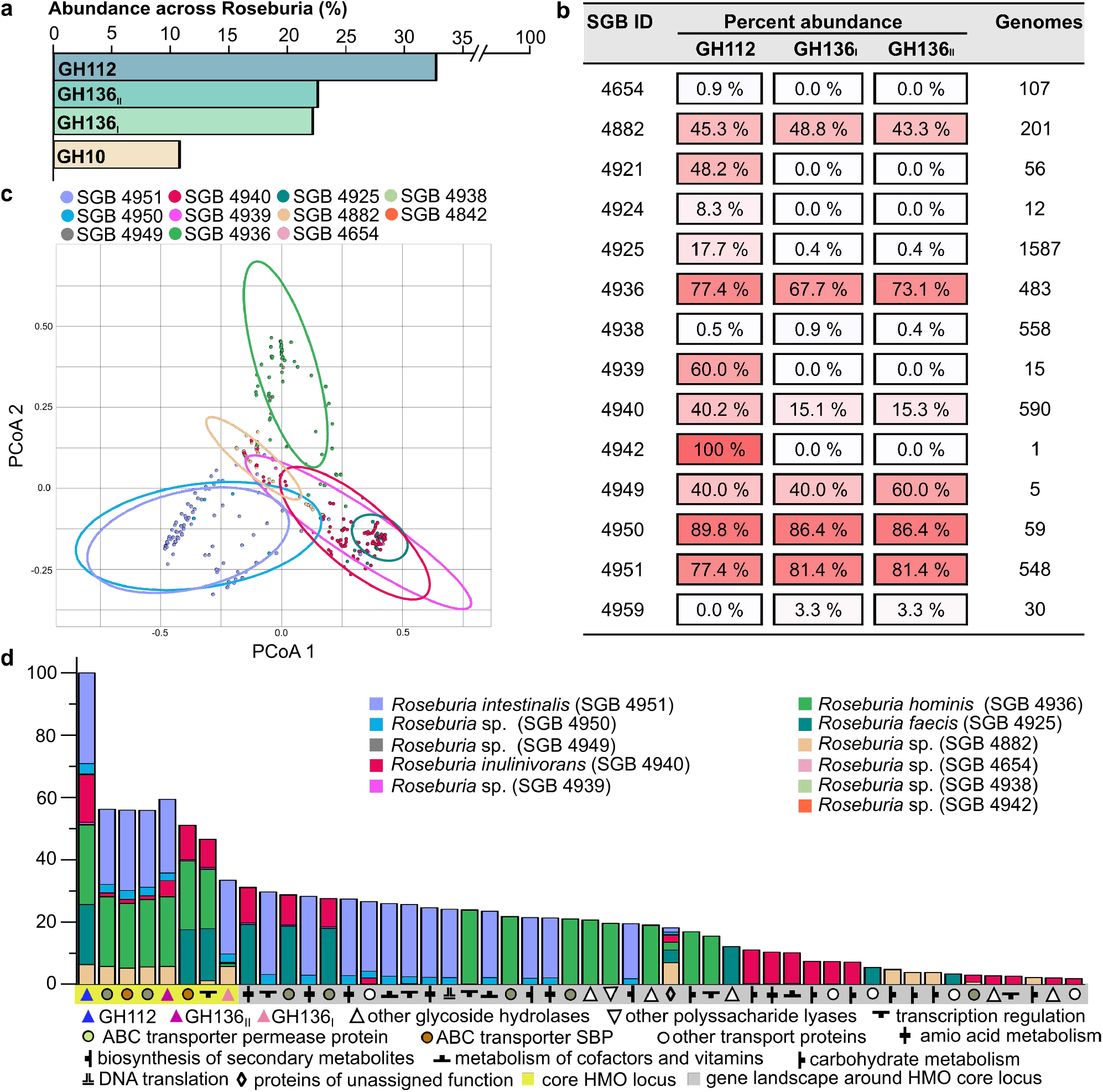
The conservation and structure of HMO utilization loci in *Roseburia.* **(a)** Global abundance of GH112, GH136_I_, GH136_II_ and GH10 xylanase genes in 4599 *Roseburia* genomes. **(b)** Heatmap showing the segregation of GH112 containing genomes from (a) into different species-level genome bins (SGBs) and the corresponding relative abundance patterns of HMO utilization proteins within each SGB. **(c)** Principal component analysis of 1397 *Roseburia* gene landscapes defined stringently based on >70% identity to the GH112 with any of the 5 references *Roseburia* genomes (Supplementary Fig. 1) and including 10 proteins up and downstream of the GH112. **(d)** The composition of gene landscapes defined in (c) viewed as the occurrence frequency relative to GH112 genes. The gene landscape analyses (**c-d**) were performed using an automatic annotation pipeline, which differentiated between two different ABC-transporter solute binding proteins (SBPs) in the core HMO locus. The *R. inulinivorans* GH136 sequences are likely to be under estimated due to their divergence from canonical lacto-*N*-biosidase counterparts e.g. from *Roseburia hominis* characterized in this study. The HMO utilization loci are defined by genes encoding at least one GH112, a GH136, an ABC-transporter, and a transcriptional regulator.

## Discussion

Perturbation of the early life HGM assembly is associated with life-long effects on the immune- and metabolic homeostasis of the host^9–12^. Breastfeeding is a key affector of the dynamics of the microbiota during infancy. Weaning marks a dramatic transition towards an adult-like structure of the HGM, which matures at the age of 2-3 and exhibits high resilience throughout adulthood^7,8,22^.

The critical window that precedes the maturation of the microbiota offers a unique opportunity for therapeutic interventions to address aberrant HGM states and thereby to prevent dysbiosis-related chronic disorders. To date, insight into the compositional signatures that characterize the assembly of the microbiota during infancy^6–8^ is available, but the underpinning mechanisms, especially during weaning, remain largely unknown. Here, we describe previously unknown pathways that confer the growth of butyrate producing Clostridiales on distinct HMO motifs and related oligomers from host glyco-conjugates. These pathways correlate to the early colonization by Clostridiales associated with the healthy HGM and with the protection from metabolic and inflammatory disorders as well as colorectal cancer^24–26,33^.

### The protein apparatus that confers the metabolism of HMO motifs and related glycoconjugate oligomers in butyrate producing Firmicutes of the HGM

We uniquely demonstrate that key butyrate producing *Roseburia* and *Eubacterium* spp. grow on complex HMOs purified from mother’s milk and on defined HMO motifs (Fig. 1a-c). Proteomic analyses revealed two highly upregulated genetic loci that encode distant homologs to a lacto-*N*-biosidase from *B. longum* ^28,29^, GNB/LNB phosphorylases and ABC transporters in *R. hominis* and *R. inulinivorans,* (Fig. 1f-g and Supplementary Fig. 1). Our analyses (Fig. 1 and 2, Supplementary Fig. 6, Table 1, Supplementary Tables 2, 3, 4 and 5) established that the locus of *R. hominis* supports growth on the HMO motifs LNT and LNB, whereas the *R. inulinivorans* locus confers growth on more complex HMOs, e.g. single and double fucosylated versions of LNT (Fig. 2a and Supplementary Fig. 6a-d). This specialization on different, but partially overlapping, HMOs and related Lewis a and Lewis b antigen oligomers from glyco-lipids or glyco-proteins creates differential competitive catabolic niches. This specialization is evident from the divergence of the GH136 specificities. Thus, *Rh*Lnb136 and *Er*Lnb136 are lacto-*N*-biosidases, whereas *Ri*Le^a/b^136 displays an unprecedented specificity that requires a Fuc-α-(1→4)-GlcNAc at the proximal glycone subsite (subsite −1) and accommodates additional fucosylation at the −2, and +2 subsites (Fig. 2, Supplementary Fig. 6a and Supplementary Table 4). The preference to fucosylation is consistent with an open active site effectuated by shortening of loops, (*Er*Lnb136: Loop 1 AA 330-341, Loop 2 AA 520-543, Supplementary Fig. 9c), which allows the accommodation of bulky fucosylated substrates. Remarkably, the GH136_I_ subunits (or domains in *Er*GH136-like enzymes) are co-evolved with the GH136_II_ counterparts that possess the catalytic residues (Supplementary Fig. 9d).

Our stability (Supplementary Fig. 8c), structural (Fig. 3 and Supplementary Fig. 9), biochemical (Supplementary Fig. 8, Supplementary Table 4) and phylogenetic analyses (Supplementary Fig. 9d) affirm the crucial role of the GH136_I_ domain in the functionality of GH136 enzymes and provide the first insight into the association of the two GH136 domains. The sequence conservation of GH136_I_ and GH136_II_ was mapped on the structure of *Er*Lnb136. Strikingly, highly conserved patches were identified across both domains (Supplementary Fig. 9e). Particularly, parts of the α4-α5 loop and of the α5 helix in *Er*Lnb136_I_ that pack extensively onto *Er*Lnb136_II_ display globally conserved residues, together with the complementary co-conserved regions of *Er*Lnb136_II_ (Supplementary Fig. 9e). Moreover, the surface of *Er*Lnb136_I_ is positively charged and apolar at the interface with *Er*Lnb136_II_, which is notably different from the negative potential on the surface of the rest of the enzyme (Supplementary Fig. 9f) and complementary to the interface surface of *Er*Lnb136_II_. These results highlight the co-evolution of GH136 subunits or domains.

ABC transporters are a determinant of uptake selectivity and competitiveness in both bifidobacteria^17,34,35^ and *Roseburia intestinalis*^32^. The two SBPs of the ABC importers located in the HMO loci of *R. hominis* and *R. inulinivorans* were within the top 5 upregulated proteins in each proteome in response to HMO utilization (Fig. 1), underscoring the critical role of oligosaccharide transport in the competitive gut niche. The preferences of the SBPs and GHs encoded by these loci appear aligned to confer efficient uptake and subsequent catabolism of preferred substrates (Fig. 2 and Supplementary Fig. 6, Table 1). The LNB/GNB phosphorylases of GH112 are also conserved in the HMO loci (Supplementary Fig. 1). *R. inulinivorans* possesses additional CAZymes, notably different fucosidases for degradation of internalized fucosylated-oligomers (Supplementary Fig. 1 and 6b-d). Based on the proteomic analyses and the biochemical data, we propose a model for the two distinct routes for uptake and depolymerisation of HMOs in *Roseburia* and *Eubacterium* (Fig. 5 and Supplementary Fig. 2).

**Fig. 5:**
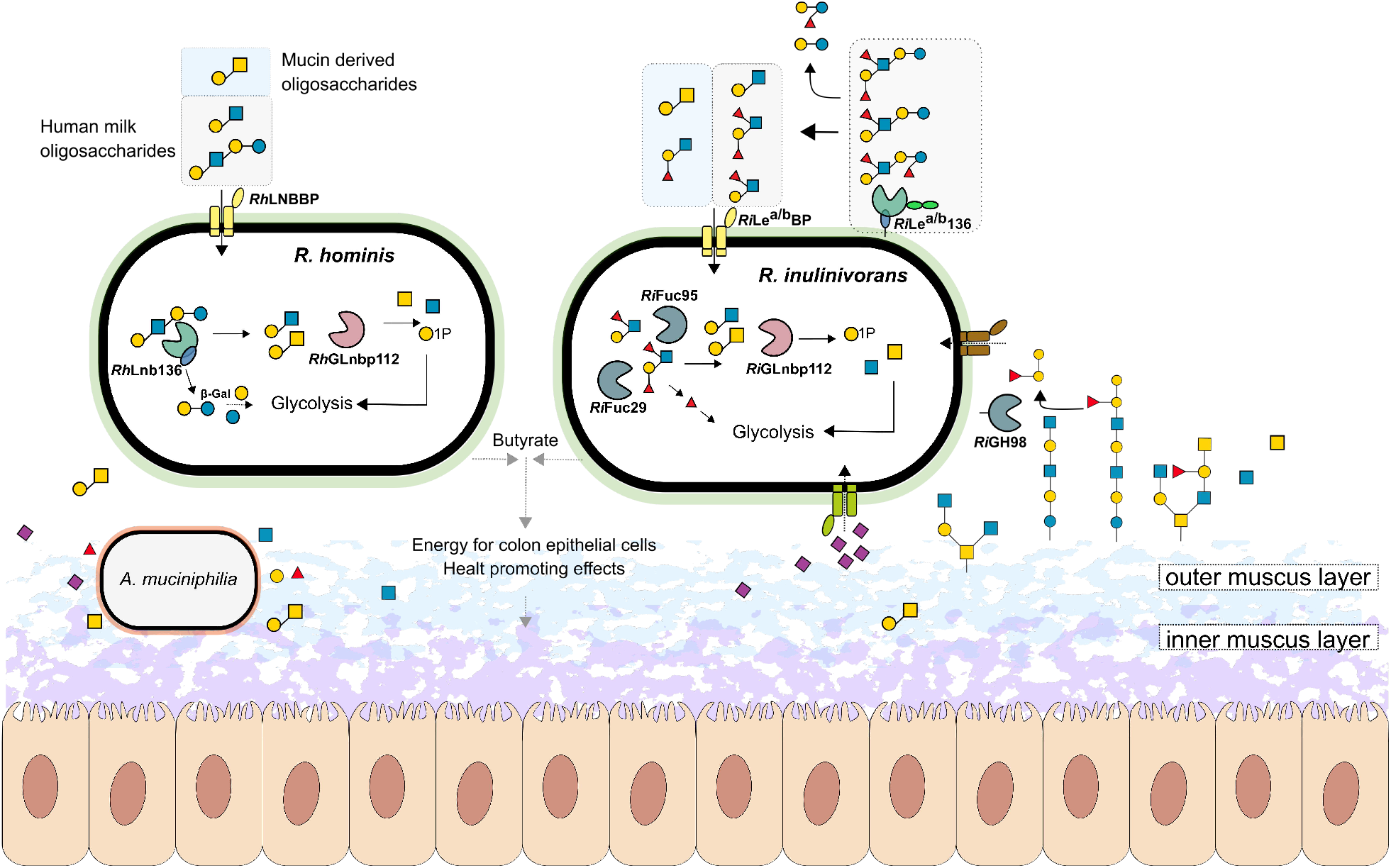
Model for HMOs and related host glycan utilization by *Roseburia* and other Lachnospiraceae. In *R. hominis* the HMOs blocks LNT and LNB and the mucin derived GNB are captured by *Rh*LNBBP for uptake into the cytoplasm and LNT is subsequently hydrolyzed to LNB. Both LNB and GNB are phosphorolyzed by *Rh*GLnbp112 into α-D-galactose-1-phosphate and the corresponding *N*-acetylhexosamines GlcNAc and GalNAc, respectively. Lactose is likely hydrolysed by a canonical β-galactosidase. *R. inulinivorans* specializes on different HMO blocks and structurally overlapping oligomers from glyco-lipids or glyco-proteins. Initial hydrolysis of HMOs or *O*-glycans from glyco-lipids/proteins occurs at the outer cell surface of *R. inulinivorans* by the activity of *Ri*Le^a/b^136, which has two C-terminal putative galactose-binding domains. The capture and import of degradation products is mediated by *Ri*Le^a/b^BP and the associated ABC transporter. In the cytoplasm, fucosyl decorations are remove by the concerted activity of *Ri*Fuc95 and *Ri*Fuc29 before *Rh*GLnbp112 phosphorolyzes the resulting LNB or imported GNB into monosaccharides, as described in *R. hominis.* The galactose and galactose-1-phosphate products are converted via the Leloir pathway to glucose-6-phosphate and *N*-acetylhexosamine sugars are converted to GlcNAc-6-phosphate before entering glycolysis. The pyruvate generated from glycolysis is partly converted to butyrate^45^. To make the model more appropriate *Roseburia* is presented in its ecological niche, the outer mucus layer^46^, together with *Akkermansia muciniphilia* as model mucin degrader. The ability of *R. inulinivorans* to cross-feed on sialic acid and to directly access host glycans is illustrated by the presence of the sialic acid uptake and catabolism machinery and by *Ri*GH98, cleaving β-(1→4)-linked blood group A and B oligosaccharides from mucin and glyco-lipids/proteins. Black solid arrows show enzymatic steps established or confirmed in this study. Black dotted arrows indicate steps based on literature. Grey dotted arrows indicate butyrate production by *R. hominis* and *R. inulinivorans* from mucin in co-culture with *A. muciniphilia.* The glycan structure key is the same as in Fig. 1.

### Conserved HMO utilization loci correlate to early colonization and resilience of Clostridiales from *Roseburia* and *Eubacterium*

Butyrate producing bacteria of the *Roseburia-Eubacterium* group (Clostridiales order) are early colonizers of the infant gut^6,8,36^ and are prevalent members of the adult HGM^22,23^.

The origin of this taxonomic group is enigmatic, but their presence in the human milk microbiome has been reported^37,38^. Orthogonal transfer from mothers based on the identification of the same *Roseburia* strains in mothers faeces, milk and the infant guts^39^ has also been proposed. *R. intestinalis* type strains have been isolated from infant faeces^40^, hinting the presence of this taxon before full transition to solid food. The pathways we have elucidated explain, at least partially, the association between HMO utilization and the early colonization of butyrate producing *Roseburia* and *Eubacterium*.

We have previously shown that the abundance of distinct bifidobacteria in guts of breast-fed infants is strongly correlated to efficient ABC transporters that capture the 2’- and 3’-fucosyl-lactose HMOs with high affinity (*K*_D_≈5 μM)^17^. The strains possessing these genes, e.g. from *Bifidobacterium longum* subspecies *infantis,* are not detected after weaning, as opposed to counterparts adept at utilizing plant-derived glycans. By contrast, the same *Clostridium* group XIVa strains that possess plant glycan utilization pathways^32,41,42^ retain HMO catabolic pathways. The simultaneous growth of *R. hominis* on LNT and the plant derived xylotetraose (Supplementary Fig. 5) demonstrates this catabolic plasticity, which likely confers competitive advantages during weaning.

The loci that target HMOs also mediate cross-feeding on mucin or other glyco-conjugate oligomers, e.g. GNB from mucin and blood antigen structures, both captured efficiently by *Roseburia* transport proteins (Table 1). This is consistent with the significant butyrate production measured in co-cultures of *Roseburia* and *A. muciniphila*^30^ (Supplementary Fig. 7c) and the upregulation of GH136-containing loci in the mucin co-culture and HMO monocultures (Fig 1 and Supplementary Fig. 7g). *R. inulinivorans* possesses an extensive mucolytic machinery revealed by the upregulation a fucose, a sialic acid (Supplementary Fig. 3) and a blood group A and B-locus (Supplementary Fig. 7h-j, Supplementary Table 9) that allows the release of β-(1→4)-linked blood group oligomers found in mucin and glyco-lipids on the surfaces of enterocytes^43,44^. This ability to access carbohydrates from mucin and host glyco-conjugates supports growth during periods of nutritional perturbations, which may increase the resilience of this taxonomic group.

Our metagenomics analyses establish that HMO utilization appears to be a core trait within *Roseburia,* based on the ubiquitousness of loci harboring GH112 and GH136 genes (Fig. 4). The presence of SGBs that exclusively possess GH112 genes (e.g. SGBs 4921 and 4939, Fig. 4b) suggests that distinct strains are secondary degraders that cross-feed on released simple substrates, e.g. LNB and GNB. By contrast, the balanced occurrence of GH112 and GH136 genes (Fig. 4b) offers a signature for primary degraders that are able to access more complex glycans from HMOs or host glyco-conjugates.

In conclusion, the present study sets the stage for a mechanistic understanding of the assembly of core physiologically important groups in the early life microbiota and discloses previously unknown roles of HMOs in selection of Clostridiales. Additional studies are required to further address the paramount, but poorly understood maturation of the early life microbiota.

## Material and Methods

### Resources: Chemicals and Carbohydrates

Human milk and blood antigen oligosaccharides used in this study are described in Table S1. *N*-acetylneuraminic acid (Neu5Ac), α-D-galactose-1-phosphate (Gal1P) and α-L-fucose (Fuc) were form Carbosynth and xylotetraose was from Megazyme. Galactose (Gal), Glucose (Glc), *N*-acetylglucosamine (GlcNAc), *N*-acetylgalactosamine (GalNAc) and porcine gastric mucin type III, (PGM) were from Sigma Aldrich. Bovine submaxillary mucin (BSM) was from VWR. 2-aminoanthranilic acid (2-AA) was from Nacalai Tesque and purified human milk samples were prepared from mother milk from Hvidøvre hospital (Hvidøvre, Denmark). All chemicals were of analytical grade unless otherwise stated.

### Method Details

#### Enzymatic production of LNB and GNB

LNB and GNB for growth were produced enzymatically with the GH112 galacto-*N*-biose/lacto-*N*-biose phosphorylase (EC 2.4.1.211) from *R. hominis* (*Rh*GLnbp112). In detail, 100 mM Gal1P and 300 mM corresponding *N*-acetylhexosamine (GlcNAc or GalNac) in 50 mM MES, 150 mM NaCl, pH 6.5 were incubated with 10 μM *Rh*GLnbp112 for 36 h at 30°C. After incubation, 2.5 volumes of ice-cold ethanol (99 %) were added, samples were incubated at – 20°C for 2 h and centrifuged (10.000x *g,* 30 min at 4°C) to remove the enzyme. Supernatants were up concentrated by rotary evaporation and disaccharides were desalted in ultrapure water (milliQ) using a HiPrep Desalt column (GE Healthcare, Denmark) on an Äkta avant chromatograph (GE Healthcare). Elution was monitored by measuring *A*_235 nm_ and pooled fractions were freeze dried. Further purification was accomplished by high-performance liquid chromatography (HPLC) (UltiMate 3000, Dionex) using a TSKgel^®^ Amide 80 column (4.6 x 250 mm) and a TSKgel^®^ Amide 80 guard column (4.6 x 10 mm) (VWR) by loading LNB or GNB dissolved in the mobile phase (75% (v/v) acetonitrile, ACN) and an isocratic elution at 1 mL min^-1^. Purity of collected fractions (2 mL) was analyzed by thin layer chromatography (TLC) using 5 mM standards of GalNAc, GlcNAc, Gal1P and LNB/GNB. Fractions containing pure LNB/GNB were pooled, ACN was removed by speed vacuum evaporation and samples were lyophilized until further use.

#### Purification of human milk oligosaccharides

Human milk oligosaccharides (HMOs) were purified from pooled human milk samples as previously described^47,48^. Milk fat was separated by centrifugation (10.000x *g,* 30 min at 4°C) and proteins were removed by ethanol precipitation (as above). The supernatant was up concentrated by rotary evaporation, buffered with 2 volumes 100 mM MES, 300 mM NaCl, pH 6.5 and lactose was digested with β-galactosidase from *Kluyvermomyces lactis* (Sigma Aldrich) (20 U mL^-1^, 3 h at 37°C). The enzyme was precipitated with ethanol (as before) and the supernatant was concentrated by rotary evaporation. Residual lactose and monosaccharides were removed by solid-phase extraction (SPE) using 12 mL graphitized Supelclean™ ENVI-Carb^TM^ columns (Supelco) with a bed weight of 1 g. For SPE, columns were activated with 80% (v/v) ACN containing 0.05% (w/v) formic acid (FA) and equilibrated with buffer A (with 4% (v/v) ACN, 0.05% (w/v) FA), which was also used to dilute the samples prior to loading. After sample loading, the columns were washed (6 column volumes of buffer A) to remove lactose and monosaccharides before oligosaccharides were eluted with 40% (v/v) ACN, 0.05% (w/v) FA. Eluted oligosaccharides were concentrated in a speed vacuum concentrator, freeze-dried and dissolved in milliQ prior to usage.

Purity of HMOs was verified by high performance anion exchange chromatography with pulsed amperometric detection (HPAEC-PAD) on an ICS-5000 (Dionex) system with a 3 × 250 mm CarboPac PA200 column (Theromofisher), a 3 × 50 mm CarboPac guard column (Theromofisher) and 10 μL injections. HMOs were eluted with a stepwise linear gradient of sodium acetate: 0-7.5 min of 0-50 mM, 7.5-25 min of 50-150 mM and 25-35 min of 150-400 mM, at a flow rate of 0.35 mL min^-1^ and a mobile phase of constant 0.1 mM NaOH. Standards (0.01-0.5 mM) of lactose, galactose and glucose in milliQ were used to quantify these residual sugars as described above. The analysis was performed in triplicates and the residual content of these sugars was <2 % (w/w) of the purified HMO mixture.

#### Isolation and purification of porcine mucins

The commercial porcine gastric mucin (PGM) was further purified^49^. In short, 20 g PGM was stirred for 20 h at 25°C in 20 mM phosphate buffer, 100 mM NaCl, pH 7.8 (adjusted to pH 7.2 after the first 2 hours using 2 M NaOH). Insoluble residues were removed by centrifugation (10,000x *g,* 30 min at 4°C) and soluble mucin was precipitated by the addition of 3 volumes of ice cold ethanol (99%) and incubation for 18 h at 4°C. Precipitated mucin was dialyzed 5 times against 200 volumes milliQ for 16 h at 4°C, using a 50 kDa molecular weight cut off membrane (Spectra, VWR) and afterwards freeze dried.

Porcine colonic mucin was isolated from five fresh pig colons from the slaughterhouse of Danish Crown (Horsens, Denmark). Pig colons were processed at site and immediately placed on dry ice to ensure quick cooling during transport. Colons were opened longitudinally and content was removed mechanically and by washing with ice cold 0.9% (w/v) NaCl until no digesta was visible. Cleaned luminal surface was quickly dried with absorptive paper and the mucosa was scraped off with a blunt metal spatula and subsequently transferred into a pre-cooled glass beaker whereby visible fat was removed and discarded. Mucin was then purified as previously described^50^. Isolated mucin was immersed in 10 volumes extraction buffer (10 mM sodium phosphate buffer, 6 M guanidine hydrochloride (GuHCl), 5 mM Ethylenediaminetetraacetic acid (EDTA), 5 mM *N*-ethylmaleimide, pH 6.5) and gently stirred overnight at 4°C. Soluble impurities and floating fat were separated by centrifugation (10,000x *g,* 30 min at 4°C), pelleted mucin was dissolved in 10 volumes extraction buffer and incubated for 3 h at room temperature again. Soluble impurities were removed by centrifugation as described before. Short incubation (3 h) extraction steps were repeated 7 times until the supernatant was clear for at least two repeated extractions. Afterwards insoluble mucin was solubilized by reduction in 0.1 M Tris, 6 M GuHCl, 5mM EDTA, 25 mM dithiotreitol (DTT) pH 8, for 5 h at 37°C and subsequent alkylation through the addition of 65 mM iodoacetamide and incubation in the dark for 18 h at 4°C. Soluble mucin was dialyzed 6 times against 200 volumes milliQ using a 50 kDa MWCO dialysis bag for 6 h at 4°C and freeze dried.

#### Cloning, expression and purification of proteins

Open reading frames encoding proteins from *R. hominis* DSM 16839, *R. inulinivorans* DSM 16841 and *E. ramulus* DSM 15684 were cloned without signal peptide or transmembrane domain from genomic DNA using In-Fusion cloning (Takara) and the primers in Table S8 into the EcoRI and Ncol restriction sites of the corresponding plasmids, to encode proteins with either a cleavable Nor C-terminal His6 tag. The pETM 11 plasmid was used (from G. Stier, EMBL, Center for Biochemistry, Heidelberg, Germany)^51^, except for *RHOM_04110* (*Rh*Lnb136_I_) and *ROSEIMA2I94_01899* (*Ri*Le^a/b^136_I_) which were cloned into pET15b (Novagen). Recombinant proteins were expressed in *E. coli* BL21 *ΔlacZ* (DE3)/pRARE2 (a kind gift from Prof. Takane Katayama, Kyoto University, Kyoto, Japan) and purified following standard protocols using His-affinity and size-exclusion chromatography. Mutants of *E. ramulus HMPREF0373_02965* (*Er*Lnb136) were constructed using QuickChange II Site-Directed Mutagenesis (Agilent) with pETM11_ *HMPREF0373_O2965* as template. Primers used for site-directed mutagenesis are listed in Table S8 and mutants were produced as described above. l-Selenomethionine labelled protein expression of *Er*Lnb136 was performed by introducing the corresponding plasmid into *E. coli* B834 (DE3) and culturing the transformed cells in a synthetic M9 based medium of the SelenoMet labelling Kit (Molecular Dimensions) supplemented either with L-methionine or l-Selenomethionine (both to 50 μg mL^-1^). The l-SeMet labelled protein was purified as described above.

#### Growth experiments and single strain proteomics analysis

*R. hominis* DSM 16839, *R. inulinivorans* DSM 16841, *E. ramulus* DSM 15684 and *E. ramulus* DSM 16296 were grown anaerobically at 37°C using a Whitley DG250 Anaerobic Workstations (Don Whitley Scientific). *R. hominis* and *R. inulinivorans* were propagated in YCFA medium^40^ while for *E. ramulus* strains CFA medium (modified YCFA medium lacking yeast extract to minimize *E. ramulus* growth on yeast extract) was used. Growth media were supplemented with 0.5% (w/v) carbohydrates sterilized by filtration (soluble carbohydrates, 0.45 μm filters) or autoclaving (mucins, 15 min at 121°C) and cultures were performed in at least biological triplicates unless otherwise indicated. Bacterial growth was monitored by measuring *OD*_600 nm_ and pH (for co-culture experiments). For growth experiments performed in microtiterplates, a Tecan Infinite F50 microplate reader (Tecan Group Ltd) located in the anaerobic workstation was used and growth was followed by measuring *OD*_595 nm_.

For differential proteome analyses, *R. hominis* and *R. inulinivorans* were grown in 200 μL YCFA (1.5 mL Eppendorf tubes) to mid-late exponential phase (*OD*_600_ ~0.5-0.8) in four biological replicates. For *R. hominis* YCFA was supplemented with 0.5% (w/v) LNT or glucose and for *R. inulinivorans* 1% (w/v) HMOs or glucose was used as carbon source. Cells were harvested by centrifugation (5.000x *g,* 5 min at 4°C), washed twice with ice cold 0.9% (w/v) NaCl, resuspended in 20 μL lysis buffer (50 mM HEPES, 6 M GuHCl, 10 mM Tris(2-carboxyethyl)phosphine hydrochloride (TCEP), 40mM 2-chloroacetamide (CAA) pH 8.5) and stored at −80°C for proteomics analysis.

#### Co-culture cross-feeding experiment proteomics analyses

*R. hominis*, *R. inulinivorans* and *A. muciniphila* DSM 22959 were grown in 10 mL YCFA to mid-late exponential phase (*OD*_600_ ~0.6-0.7). From these pre-cultures, equal amounts of cells (*OD*_600_) were used to inoculate 30 mL fresh YCFA medium with 1% (w/v) of a mucin mixture (0.6% (w/w) PGM, 0.2% (w/w) PCM, 0.2% (w/w) BSM) or 1% (w/v) glucose to a start *OD*_600_ ~0.01. All cultures were performed in four biological replicates and growth was followed (*OD*_600_ and pH) at 0, 6, 8, 12, 16, 24, and 48 h. Samples (2 mL) were collected for proteomics analyses after 16 h and for SCFA quantification after 24 and 48 h. Samples were immediately cooled on ice and cells were harvested by centrifugation (5000x *g,* 10 min at 4°C). For proteomics, cell pellets were washed twice with ice cold 0.9% (w/v) NaCl, resuspended in 60 μL lysis buffer and stored at −80°C until proteomics analysis. Collected culture supernatants for SCFA quantification were sterile filtrated (0.45 μm filters) and stored at −80°C for further analysis.

#### Protein extraction and sample preparation for mass spectrometry

Samples were processed as described elsewhere^52,53^. Cells were lysed by boiling (5 min 95°C) followed by bead beating (3mm beads, 30 Hz for 1 min) (TissueLyser II, Qiagen) and sonication bath (3×10 sec at 4°C) (Bioruptor, Diagenode). Lysates were centrifuged (14.000x *g*, 10 min at 4°C) and soluble protein concentrations were determined by a Bradford assay (Thermo Fisher Scientific). For digestion, 20 μg protein were diluted 1:3 with 50 mM HEPES, 10% (v/v) ACN, pH 8.5 and incubated with LysC (MS grade, Wako) in a ratio of 1:50 (LysC:protein) for 4 h at 37°C. Subsequently, samples were diluted to 1:10 with 50 mM HEPES, 10% (v/v) ACN, pH 8.5 and further digested with trypsin (MS grade, Promega) in a ratio of 1:100 for 18 h at 37°C. Next, samples were diluted 1:1 with 2% (w/v) trifluoroacetic acid (TFA) to quench enzymatic activity and peptides were processed for mass spectrometry using in house packed stage tips^54^ as described below.

Peptides from single strain cultures were desalted using 3 discs of C18 resin packed into a 200 μL tip and activated by successive loading of 40 μL of MeOH and 40 μL of 80% (v/v) ACN, 0.1% (w/v) FA by centrifugation at 1800x *g* and equilibrated twice with 40 μL of 3% (v/v) ACN, 1% (w/v) FA before samples were loaded in steps of 50 μL. After loading, tips were washed three times with 100 μL 0.1% (w/v) TFA and peptides were eluted in two steps with 40 μL each of 40% (v/v) ACN, 0.1% (w/v) FA into a 0.5 mL Eppendorf LoBind tube. Peptides derived from co cultures were desalted and fractionated using strong cation exchange (SCX) chromatography filter pugs (3M Empore). Per sample, 6 SCX discs were packed into a 200 μL tip and tips were activated and equilibrated by loading 80 μL (v/v) of ACN and then 80 μL of 0.2% (w/v) TFA. Samples were applied in 50 μL steps and tips were washed twice with 600 μL 0.2% (w/v) TFA. Subsequently peptides were stepwise eluted in 3 fractions with 60 μL of 125 mM NH4OAc, 20% (v/v) ACN, 0.5% (w/v) FA, then with 60 μL of 225 mM NH_4_OAc, 20% (v/v) ACN, 0.5% (w/v) FA and lastly with 5% (v/v) NH_4_OH, 80 % (v/v) ACN into 0.5 mL Eppendorf LoBind tubes. Eluted peptides were dried in an Eppendorf Speedvac (3 h at 60°C) and reconstituted in 2% (v/v) ACN, 1% (w/v) TFA prior to mass spectrometry (MS) analysis.

#### LC-MS/MS

Peptides from biological triplicates of each culture condition were loaded on the mass spectrometer by reverse phase chromatography through an inline 50 cm C18 column (Thermo EasySpray ES803) connected to a 2 cm long C18 trap column (Thermo Fisher 164705) using a Thermo EasyLc 1000 HPLC system. Peptides were eluted with a gradient of 4.8–48 % (v/v) ACN, 0.1% (w/v) FA at 250 nL min^-1^ over 260 min (samples from single strain cultures) or 140 min (SCX fractionated samples from co cultures) and analysed on a Q-Exactive instrument (Thermo Fisher Scientific) run in a data-dependent manner using a “Top 10” method. Full MS spectra were collected at 70,000 resolution, with an AGC target set to 3×10^6^ ions or maximum injection time of 20 ms. Peptides were fragmented via higher-energy collision dissociation (normalized collision energy=25). The intensity threshold was set to 1.7×10^6^, dynamic exclusion to 60 s and ions with a charge state <2 or unknown species were excluded. MS/MS spectra were acquired at a resolution of 17,500, with an AGC target value of 1×10^6^ ions or a maximum injection time of 60 ms. The scan range was limited from 300–1750 m/z.

#### Protein identification and Label-free quantification and relative abundance in co-culture communities

Proteome Discoverer versions 2.2 & 2.3 were used to process and analyze the raw MS data files and label free quantification was enabled in the processing and consensus steps. The spectra from single strains proteomics were matched against the proteome database of *R. hominis* DSM 16839 (ID: UP000008178) or *R. inulinivorans* DSM 16841 (ID: UP000003561) respectively, as obtained from Uniprot. The spectra from co-culture experiments were searched against a constructed database consisting of the reference proteomes of the two *Roseburia strains* (as above) and *A. muciniphila* DSM 22959 (ID: UP000001031). For spectral searches, oxidation (M), deamidation (N, Q) and N-terminal acetylation were specified as dynamic modifications and cysteine carbamidomethylation was set as a static modification. Obtained results were filtered to a 1% FDR and protein quantitation was done by using the built-in Minora Feature Detector. For analysis of the label-free quantification data, proteins were considered present if at least 2 unique peptides (as defined in Proteome Discoverer) were identified and proteins had to be identified in at least 2 out of the 3 samples analyzed per culture condition with high confidence.

Relative bacterial abundance in co-cultures was estimated based on strain unique peptides identified with Unipept version 4.0^55^. To exclude peptides shared between closely related strains from the analyses, all peptide sequences quantified via Proteome Discoverer were imported into the Unipept web server and analyzed with the settings “Equate I and L” and “Advanced missed cleavage handling” activated. The normalized sum of intensities of the resulting taxonomically distinctive peptides was then used for assessing relative abundances of each strain.

#### Butyrate quantification

Butyrate in culture supernatants was quantified by HPLC coupled to a refracting index detector (RID) and diode array detector (DAD) on an Agilent HP 1100 system (Agilent). Standards of butyric acid (0.09-50 mM) were prepared in 5 mM H2SO4 for peak identification and quantification. Samples from 4 biological replicates were analysed by injecting 20 μL of standard or filtrated (0.45 μM filter) culture supernatant on a 7.8 x 300 mm Aminex HPX-87H column (Biorad) combined with a 4.6 x 30 mm Cation H guard column (Biorad). Elution of was performed with a constant flow rate of 0.6 mL min^-1^ and a mobile phase of 5 mM H2SO4. Standards were analysed as above in technical triplicates.

#### Oligosaccharide uptake preference of *R. hominis*

*R. hominis* was grown anaerobically in 250 μL YCFA medium with 0.5% (w/v) of an equal mixture of xylotetraose and LNT in biological triplicates. Samples (20 μL) were taken after 0, 3.5, 5.5, 6.5, 8, 9.5 and 24h, diluted 10-fold in ice cold 100 mM NaOH and centrifuged (10 min at 5000x *g* at 4°C) before supernatants were stored at −20°C until the HPAEC-PAD analysis. Standards of 0.5 mM xylotetraose and LNT were prepared in 100 mM NaOH and used to identify corresponding peaks in the chromatograms. Samples or standard were injected (2 μL injections) on a 4 × 250 mm CarboPac PA10 column with a 4 × 50 mm CarboPac guard column and eluted isocratically (0.750 mL min^-1^, 100 mM NaOH, 10mM NaOAc). The analysis was performed from a biological triplicate and standards were analyzed in technical duplicates.

#### Enzyme activity assays

Enzymatic activity assays were carried out in 50 mM MES, 150 mM NaCl, 0.005% (v/v) Triton X-100, pH 6.5 standard assay buffer and in triplicates unless otherwise stated.

Hydrolysis kinetics and specific activities of the GH136 lacto-*N*-biosidases were measured using a coupled enzymatic assay to monitor lactose release. The lactose was hydrolyzed with a β-galactosidase (used above) and the resulting glucose was oxidized with a glucose oxidase (Sigma Aldrich) concomitant with the production of H_2_O_2_ measured by coupling to horseradish peroxidase (Sigma Aldrich) oxidation of 4-aminoantipyrine and 3,5-dichloro-2-hydroxybensensulfonic acid. Reactions were prepared in 96-well microtiter plates to a final volume of 150 μL, containing substrate, lacto-*N*-biosidase, β-galactosidase (150U mL^-1^), glucose oxidase (150U mL^-1^), horseradish peroxidase (150U mL^-1^), 10 mM 3,5-dichloro-2-hydroxybensensulfonic acid, 1 mM 4-aminoantipyrine in standard assay buffer. Reactions were performed at 37°C and *A*_515_ nM was measured in 5 sec intervals for 30 min. Blanks were prepared by substituting lacto-*N*-biosidase with standard assay buffer in the reaction mixture and a lactose standard (3 μM-500 μM) was used for the quantification.

Hydrolysis kinetics of *Rh*Lnb136 (40 nM) and *Er*Lnb136 (10 nM) towards LNT (0.2–5 mM for *Rh*Lnb136 and 0.1-2.5 mM for *Er*Lnb136) were determined as described above. The kinetic parameters *K_M_* and *k*_cat_, were calculated by fitting the Michaelis-Menten equation to the initial rate data using OriginPro 2018b (OriginLab). Lacto-*N*-biosidase specific activity of *Ri*Le^a/b^136 (1.2 μM) was measured as described above using 3.5 mM LNT. The specific activity was expressed in units (U) mg^-1^ enzyme, where a unit is defined as the amount of enzyme that releases 1 μmol lactose min^-1^ quantified as above.

Specific activities of *Rh*GLnbp112 and *Ri*GLnbp112 towards LNB and GNB were assayed 50 mM sodium phosphate buffer, 150 mM NaCl, 0.005% (v/v) Triton X-100, pH 6.5. Reactions (150 μL) were incubated for 10 min at 37°C with 20 nM enzyme and 2 mM substrate. Aliquots of 15 μL were removed every minute and quenched in 135 μL 0.2 M NaOH. Standards of Gal1P (5 mM–0.02 mM) were prepared in 0.2 M NaOH and were used to quantify the concentrations of released Gal1P in the quenched reaction samples. Both, quenched reactions and standards were examined by HPAEC-PAD using a 3 × 250 mm CarboPac PA200 column (Theromofisher) in combination with a 3 × 50 mm CarboPac guard column (Theromofisher) and 10 μL injections. Elution was performed with a flow of 0.350 mL min^-1^ and a mobile phase of 150 mM NaOH and 60 mM sodium acetate. The specific activity was expressed in U mg^-1^ enzyme, where a U is defined as the amount of enzyme that releases 1 μmoL Gal 1P min^-1^. The analysis was performed in technical triplicates.

#### Enzyme product profiles

Enzyme assays were performed at 37°C for 16 h in standard assay buffer or in the phosphate version (instead of MES) for GH112 enzymes. Degradation products were analyzed by thin layer chromatography (TLC) and or Matrix-assisted laser desorption/ionization time of flight mass spectroscopy (MALDI-TOF/MS) as described below.

#### Thin layer chromatography

The TLC was performed by spotting 2 μL of enzymatic reaction on a silica gel 60 F454 plate (Merck), the separation was carried out in butanol: ethanol: milliQ water (5:3:2) (v/v) as mobile phase and sugars were visualized with 5-methylresorcinol:ethanol:sulfuric acid (2:80:10) (v/v) and heat treatment except for *Ri*Le^a/b^136. The TLC for the latter enzyme was performed in butanol:acetic acid: milliQ (2:1:1)(v/v) and developed with diphenylamine-phosphoric acid reagent^56^.

#### MALDI-TOF/MS

MALDI-TOF/MS analysis of *Ri*Le^a/b^136 was accordind to^57^, following permethylation of oligosaccharides^58^. Permethylated sugars were dried, mixed with 2,5-dihydroxybenzoic acid, and spotted onto the MALDI plate. For MALFI-TOF/MS analyses, a Bruker Autoflex III smartbeam in positive ion mode was used. Degradation products of *Rh*Lnb136 and *Er*Lnb136 were analyzed without initial permethylation of oligosaccharides using 2,5-dihydroxybenzoic acid as matrix and an Ultraflex II TOF/TOF (Bruker Daltonics) instrument operated in positive ion linear mode. Peak analysis of mass spectra was performed using Flexanalysis Version 3.3 (Bruker Daltonics).

#### LC-MS^2^ of *O*-glycan derived oligosaccharides

A homogenous preparation of porcine gastric mucin, PGM (Sigma), carrying blood group A, was used in the analysis. A total of 0.1 mg mucin per dot were immobilized by dot blotting onto an immobilon-P PVDF membranes (Immobilon P membranes, 0.45 μm, Millipore, Billerica, MA). *Ri*GH98 was added to one dot to 1.5 μM in 50 μL and incubated for 1 h and 4 h at 37°C. The reaction supernatants which contained released free oligosaccharides, were collected and purified by passage through porous graphitized carbon (PGC) particles (Thermo Scientific) packed on top of a C18 Zip-tip (Millipore). Samples were eluted with 65% (v/v) ACN in 0.5% trifluoro-acetic acid (TFA, v/v), dried, resuspended in 10 μL of milliQ, frozen at −20 °C and stored until further analysis. The residual *O*-linked glycans (on the dot) were released by reductive β-elimination by incubating the dot in 30 μL of 0.5 M NaBH_4_ in 50 mM NaOH at 50°C for 16 h followed by adding 1.5 μL glacial acetic acid to quench the reaction. The released *O*-glycans were desalted and dried as described before ^59^. The purified glycans were resuspended in 10 μL of milliQ and stored at −20°C for further analysis. Released oligosaccharides from glycosphingolipids as a model substrate carrying blood group B (B5-2 and B6-2)^60^ were prepared as described above, except for a single incubation time of 2 h.

Purified samples were analyzed by LC-MS/MS using 10 cm × 250 μm I.D. column, packed in house with PGC 5 μm particles. Glycans were eluted using a linear gradient of 0–40% ACN in 10 mM NH_4_HCO_3_ over 40 min at 10 μl min^-1^. The eluted *O*-glycans were analyzed on a LTQ mass spectrometer (Thermo Scientific) in negative-ion mode with an electrospray voltage of 3.5 kV, capillary voltage of −33.0 V and capillary temperature of 300 °C. Air was used as a sheath gas and mass ranges were defined depending on the specific structure to be analyzed. The data were processed using Xcalibur software (version 2.0.7, Thermo Scientific).

#### Oligosaccharide binding analysis

Binding of LNT, LNB, GNB, H type I triose, Le^a^ triose and Le^b^ tetraose to *Ri*Le^a/b^BP was analyzed by surface plasmon resonance (SPR; Biacore T100, GE Healthcare). *Ri*Le^a/b^BP, diluted in 10 mM NaOAc buffer pH 3.75 to 50 μg mL^-1^, was immobilized on a CM5 chip using a random amine coupling kit (GE Healthcare) to a final chip density of 3214 and 4559 response units (RU). Analysis comprised 90 s for association and 240 s for dissociation phase, respectively, at a flow rate of 30 μL min^-1^. Sensograms were recorded at 25°C in 20 mM sodium phosphate buffer, 150 mM NaCl, 0.005% (v/v) P20 (GE Healthcare), pH 6.5. Experiments were performed in duplicates (each consisting of a technical duplicate) in the range of 0.3-50 μM for LNB, 0.78-200 μM for GNB, 0.97-250 μM for Le^a^, 0.097 μM-100 μM for Le^b^ and 1.5-250 μM for blood H type I triose. To investigate ligand specify of *Ri*Le^a/b^BP, binding was further tested towards 0.5 mM LNT, LN*n*T, lactose, blood A triose, 2’FL and 3’FL. Equilibrium dissociation constants (*K*_D_) were calculated by fitting a one binding site model to steady state sensograms, using the Biacore T100 data evaluation software.

Binding of LNT, LNB, GNB, LNnT, lactose and 2’FL to *Rh*LNBBP was measured using a Microcal ITC200 calorimeter (GE Healthcare). Titrations were performed in duplicates at 25°C with *Rh*LNBBP (0.1 mM) in the sample cell and 1.5 mM ligand in 10 mM sodium phosphate buffer, pH 6.5 in the syringe. A first injection of 0.4 μL was followed by 19 injections of 2 μL ligand each, separated by 180 s. Heat of dilution was determined from buffer titrations and corrected data were analyzed using MicroCal Origin software v7.0. To determine binding thermodynamics a non-linear single binding model was fitted to the normalized integrated binding isotherms.

#### Differential scanning calorimetry (DSC)

The DSC analyses was performed at protein concentrations of 1 mg mL^-1^ in 20 mM sodium phosphate buffer, 150 mM NaCl, pH 6.5, using a Nano DSC (TA instruments). Thermograms were recorded from 10 to 90°C at a scan speed of 1°C min^-1^ using buffer as reference. Baseline corrected data were analyzed using the NanoAnalyze software (TA instruments). DSC analyses were performed in duplicates unless otherwise states.

#### Crystallization

Crystals of *Er*Lnb136 proteins were grown at 20°C using the sitting-drop vapor diffusion method, by mixing 0.5 μL of a 10 mg mL^-1^ protein solution with an equal volume of a reservoir solution. Native crystals were grown in a 20% (w/v) PEG4000, 0.1 M sodium citrate pH 5.6, and 20% isopropanol reservoir solution. SeMet-labelled crystals were grown using a reservoir solution containing 20% (w/v) PEG6000, 0.1 M Tris-HCl pH 8.5, and 1 M lithium chloride. The crystals were cryoprotected in the reservoir solution supplemented with 20% (v/v) glycerol and 25 mM LNB. The crystals were flash-cooled at 100 K (−173.15°C) in a stream of nitrogen gas. Diffraction data were collected at 100 K on beamlines at SLS X06DA (Swiss Light Source, Swiss) and Photon Factory of the High Energy Accelerator Research Organization (KEK, Tsukuba, Japan). The data were processed using HKL20 00^61^ and XDS^62^. Initial phase calculation, phase improvement, and automated model building were performed using PHENIX^63^. Manual model rebuilding and refinement was achieved using Coot^64^ and REFMAC5^65^. Because the crystal structures of SeMet-labelled and native protein were virtually the same (root mean square deviations for the Cα atoms = 0.14 Å), we used the SeMet-labelled protein structure for the descriptions in the Results and Discussion. Molecular graphics were prepared using PyMOL (Schrödinger, LLC, New York) or UCSF Chimera (University of California, San Francisco)

#### Bioinformatics

SignalP v.4.1^66^, PSORTb v3.0^67^, TMHMM v.2.0^68^ were used for prediction of signal peptides and transmembrane domains. InterPro^69^ and dbCAN2^70^ were used to analyse modular organization using default settings for Gram positive bacteria. Redundancy in biological sequence datasets was reduced using the CD-HIT server (sequence identity cut off = 0.95)^71^. Protein sequence alignments were performed using MAFFT (BLOSUM62)^72^. Phylogenetic trees were constructed using the MAFFT server, based on the neighbor-joining algorithm, and with bootstraps performed with 1000 replicates. Phylogenetic trees were visualized and tanglegrams constructed using dendroscope^73^. Coloring of protein structures according to amino acid sequence conservation was accomplished in UCSF Chimera, based on protein multiple (structural based) alignments from the PROMALS3D server^74^ and by using the in UCSF Chimera implemented AL2CO algorithm^75^. The MEME suite web server was used for amino acid sequence motif discovery and evaluation^76^. Protein structures were compared using the Dali server (http://ekhidna2.biocenter.helsinki.fi/dali/) (PMID: 27131377) and the molecular interface between *Er*Lnb136_I_ and *Er*Lnb136_II_ was analyzed (solvent inaccessible interface, Gibbs energy) via the PDBePISA server (https://www.ebi.ac.uk/pdbe/pisa/).

The abundance and distribution of HMO utilization genes encoding GH112, GH136_I_ and GH136_II_ in *Roseburia* were analyzed by a BLAST search of the corresponding DNA reference sequences from *R. intestinalis* L1-82, *R. hominis* A2-183 and *R. inulinivorans* A2-194 against a total of 4599 reconstructed *Roseburia* genomes, binned into 42 Species-level Genome Bins (SGBs) by Pasolli et al.^31^. The variability of the *Roseburia* core xylanase (GH10) was determined similarly by blasting the DNA reference sequences from *R. intestinalis* L1-82 (ROSINTL182_06494) against the same dataset.

For further analyses, initial blast hits were filtered based on a 70% identity with any of the 5 conserved *Roseburia* reference genomes. Additionally, Roseburia genomes were considered only if they have a hit with GH112 gene. The resulting 1397 genomes were assigned into the respective *Roseburia* SGBs, base on they assignation of Pasolli et al.^31^. The retrieved genomes were used to analyze the gene landscape around the GH112 gene. The RAST server^77^ was used for gene annotation. Based on the annotation and coordinates of the genes, 10 genes upstream and downstream the GH112 were selected for gene landscapes analysis.

### Quantification and Statistical Analysis

Statistical significant differences were determined using unpaired two-tailed Student’s *t*-test. Statistical parameters, including values of n and *p*-values, are reported or indicated in the figures, figure legends and the result section. The data are expressed as arithmetic means with standard deviations (SD), unless otherwise indicated.

### Data and Code availability

The mass spectrometry proteomics data have been deposited to the ProteomeXchange Consortium via the PRIDE partner repository with the dataset identifier PXD015045. The accession numbers for the atomic coordinates reported in this paper are PDB: 6KQS (Se-Met) and 6KQT (native), see also Table S6. Mucin glycomis MS/MS data are summarized in Table S9 and raw data files are available upon request.

### Material and Resource availability

Requests for resources and material should be addressed to Maher Abou Hachem

## Acknowledgement

We thank Drs Ayaka Harada and Miki Senda, and the staff of the Photon Factory and Swiss Light Source (grant numbers: 20181219 & 20181299) for the X-ray data collection. LC-MS/MS analysis of glycans was performed by the Swedish infrastructure for biological mass spectrometry (BioMS) supported by the Swedish Research Council. We also wish to thank Tina Johansen for the technical help in performing the HPLC measurements for the quantification of butyrate. Dr. Takatoshi Arakawa is thanked for managing the X-crystallography structural data. Drs. Fumihiko Sato and Kentro Ifuku are thanked for the technical support of MALDI-TOF/MS analysis. We also thank Prof. Tine Rask Licht for the use of the microplate reader for some of the growth experiments.

