## Supplemental Tables and figures for "Butyrate producing Clostridiales utilize distinct human milk oligosaccharides correlating to early colonization and prevalence in the human gut"

### Supplementary Information:

#### Supplementary Tables:

**Supplementary Table 1: Human milk and blood antigen derived oligosaccharides studied in this work**

| Compound[a] | Abbreviation | Supplier | Structure[b] |
|---|---|---|---|
| Lactose | Lac | Sigma Aldrich | 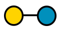 |
| 2-Fucosyllactose | 2'FL | IsoSep | 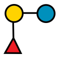 |
| 3-Fucosyllactose | 3'FL | IsoSep | 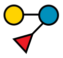 |
| Difucosyllactose | DFL | Dextra Laboratories | 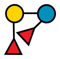 |
| Lacto-*N*-biose | LNB | Elicityl | 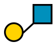 |
| Galacto-*N*-biose | GNB | Sigma Aldrich | 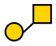 |
| Lacto-*N*-tetraose | LNT | Elicityl<br>IsoSep | 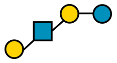 |
| Lacto-*N*-*neo*tetraose | LN*n*T | Dextra Laboratories | 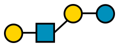 |
| Lacto-*N*-fucopentaose I | LNFP I | Dextra Laboratories | 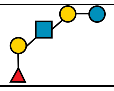 |
| Lacto-*N*-fucopentaose II | LNFP III | Hneywell Fluka | 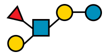 |
| Lacto-*N*-fucopentaose III | LNFP III | Carbosynth | 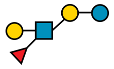 |
| Lacto-*N*-difucohexaose I | LNDFH I | Dextra Laboratories<br>Carbosynth | 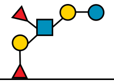 |
| Lacto-*N*-difucohexaose II | LNDFH II | Elicityl<br>Dextra Laboratories | 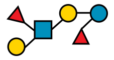 |
| Blood group antigen H triose type 1 | H triose type 1 | Elicityl | 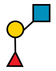 |
| Blood group antigen A triose | A triose | Elicityl | 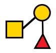 |
| Lewis A antigen triose | $Le^a$ triose | Elicityl | 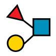 |
| Lewis B antigen tetraose | $Le^b$ tetraose | Carbosynth | 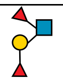 |

[a]All carbohydrates were > 95% pure unless otherwise stated.
[b]Glycan structures presentation according to Symbol Nomenclature for Glycans (SNFG) (https://www.ncbi.nlm.nih.gov/glycans/snfg.html)

**Supplementary Table 2: Binding and thermodynamic parameters of *Rh*LNBBP determined by ITC.**

| Ligand | $K_D$ (μM) | $N_0$ | $\Delta H$ (kcal mol$^{-1}$) | $-T\Delta S$ (kcal mol$^{-1}$) | $\Delta G$ (kcal mol$^{-1}$) |
|---|---|---|---|---|---|
| LNB | 2.86 ± 0.26 | 0.85 ± 0.02 | -29.25 ± 0.86 | 21.69 | -7.65 |
| GNB | 11.04 ± 0.06 | 1.25 ± 0.03 | -12.93 ± 0.42 | 6.17 | -6.75 |
| LNT | 10.30 ± 0.60 | 0.88 ± 0.01 | -19.36 ± 0.23 | 12.55 | -6.85 |
| Lactose | n.b. | | | | |
| 2’FL | n.b. | | | | |

Data are from duplicates and binding parameters are means of duplicates with standard deviation. n.b.: affinity too low to be determined. $N_0$ is the molar binding stoichiometry.

**Supplementary Table 3: Binding parameters of *Ri*Le$^{a/b}$BP determined by SPR**

| Ligand | $K_D$ (μM) | $R_{max}$ | $\chi^2$ |
|---|---|---|---|
| LNB | 6.7 ± 0.7 | 10.4 | 0.10 |
| GNB | 11 ± 0.9 | 10.4 | 0.06 |
| Le$^b$ tetraose | 1.8 ± 0.1 | 25.1 | 0.54 |
| Le$^a$ triose | 3.2 ± 0.5 | 17.3 | 0.32 |
| H triose type 1 | 11.3 ± 2.5 | 8.8 | 0.06 |
| LNT | n.b. | | |
| Blood group A antigen triose | n.b. | | |
| 2’FL | n.b. | | |
| 3’FL | n.b. | | |
| LN*n*T | n.b. | | |
| Lactose | n.b. | | |

The binding parameters are means of duplicates with standard deviation. n.b. indicates low affinity to ligand precluding determination of binding parameters. $R_{max}$ and $\chi 2$ denote the maximum binding level from the fits to a one binding site model and the statistical goodness of the fit to the same model, respectively.

**Supplementary Table 4. Kinetic parameters of *Rh*Lnb136, *Ri*Le$^{a/b}$136 and *Er*Lnb136**

| Substrate | Enzyme | $K_M$ (mM) | $k_{cat}$ ($s^{-1}$) | $k_{cat}/K_M$ ($s^{-1}$ $mM^{-1}$) | specific activity[a] (U $mg^{-1}$) |
|---|---|---|---|---|---|
| LNT | *Rh*Lnb136 | 1.45 ± 0.05 | 86 ± 1 | 59.3 | 58.5 ± 0.58 |
| LNT | *Ri*Le$^{a/b}$136 | - | - | - | 0.21 ± 0.00 |
| LNT | *Er*Lnb136 | 0.68 ± 0.07 | 160 ± 7 | 235.3 | |
| LNT | *Er*Lnb136 Y145A | n.d. | n.d. | 48 | |

[a] specific activity determined towards 3.5 mM LNT. n.d.: Lack of curvature of the Michaelis Menten plot preclude determination of kinetic parameters. Data are means of triplicates with standard deviation.

**Supplementary Table 5. Specific activities of *Rh*GLnbp112 *and Ri*GLnbp112**

| Substrate | ***Rh*GLnbp112** (U $mg^{-1}$) | ***Ri*GLnbp112** (U $mg^{-1}$) |
|---|---|---|
| LNB | 12.2 ± 0.5 | 22.6 ± 0.2 |
| GNB | 9.6 ± 0.1 | 16.9 ± 0.4 |

Data are means of triplicates with standard deviation. Specific activities determined towards 2 mM LNB or GNB.

**Supplementary Table 6: Crystallographic data collection and refinement statistics.**

| | *Er*Lnb136 Se-Met | *Er*Lnb136 Native |
|---|---|---|
| **PDB ID** | 6KQS | 6KQT |
| **Data collection**[a] | | |
| Beamline | SLS X06DA | KEK-PF BL5A |
| Wavelength (Å) | 0.978 | 1.000 |
| Space group | $P3_121$ | $P3_121$ |
| Unit cell (Å) | $a = b = 132.7$, $c = 82.5$ | $a = b = 132.3$, $c = 82.2$ |
| Resolution (Å) | 45.75–1.40 (1.42–1.40) | 50.0–2.00 (2.03–2.00) |
| $R_{merge}$ | 0.145 (1.909) | 0.233 (1.065) |
| Number of observations | 3,288,573 (155,651) | 556,706 |
| Unique reflections | 153,888 (8,031) | 56,318 (2,757) |
| Mean $I/\sigma(I)$ | 12.2 (1.7) | 13.4 (3.0) |
| CC (1/2) | 0.999 (0.728) | 0.980 (0.824) |
| Completeness (%) | 100.0 (100.0) | 100.0 (100.0) |
| Multiplicity | 20.1 (19.4) | 9.9 (9.6) |
| Anomalous completeness (%) | 100.0 (100.0) | – |
| Anomalous multiplicity | 10.2 (9.8) | – |
| **Refinement** | | |
| Resolution | 47.20–1.40 | 47.04–2.00 |
| No. of reflections | 155,798 | 53,440 |
| $R$ factor/$R_{free}$ (%) | 14.8 (17.2) | 14.3 (18.1) |
| No. of atoms | 5,920 | 5,787 |
| No. of solvents | 835 (water), 1 (glycerol) | 706 (water), 1 (triethylene glycol), 1 ($Na^+$) |
| RMSD from ideal values | | |
| Bond lengths (Å) | 0.016 | 0.011 |
| Bond angles (°) | 1.975 | 1.63 |
| Ramachandran plot (%) | | |
| Favored | 95.9 | 95.8 |
| Allowed | 4.1 | 4.2 |
| Outlier | 0 | 0 |

[a]Values in parentheses are for the highest resolution shell.

**Supplementary Table 7: Summary of structural similarity Dali server search of *Er*Lnb136.**

| Protein | Source organism | PDB (chain) | Z score | RMSD (Å) | $N_{align}$[a] | $\%_{seq}$[b] |
|---|---|---|---|---|---|---|
| ***Er*Lnb136$_{I}$ (residues 7-224)** | | | | | | |
| SurA-like putative peptidyl-prolyl cis-trans isomerase | *Helicobacter pylori* | 5EZ1 (A) | 7 | 3.2 | 70 | 19 (6) |
| Hypothetical protein LIC12922 | *Leptospira interrogans* | 3NRK (A) | 5.8 | 4.8 | 105 | 10 (5) |
| ***Er*Lnb136$_{II}$ (residues 242-663)** | | | | | | |
| LnbX (GH136) | *Bifidobacterium longum* | 5QQC (H) | 50.3 | 1.4 | 416 | 44 (43) |
| α-L-fucosidase BT_1002 (GH141) | *Bacteroides thetaiotaomicron* | 5MQP (F) | 32 | 3.1 | 312 | 16 (12) |

Data were obtained using Dali server.

[a]Number of aligned residues

[b]Sequence identity of aligned residues and the corresponding overall (global) sequence identity showed in parenthesis

**Supplementary Table 8. Primers for cloning and mutagenesis.**

| Locus tag | Accession[a] | Designation | Orientation | Sequence (5'->3') |
|---|---|---|---|---|
| RHOM_04115 | G2T0V1 | *Rh*Lnb136$_{II}$ | Forward | AGGAGATATACCATGGATGACAGGCTCATACAGGAC |
| RHOM_04115 | G2T0V1 | *Rh*Lnb136$_{II}$ | Reverse | GGTGGTGGTGCTCGAGGCCCAACGGAATAATCGTATTATCC |
| RHOM_04110 | G2T0V0 | *Rh*Lnb136$_{I}$ | Forward | AGGAGATATACCATGGATGAATCGGAAATATTGTCTGGAT |
| RHOM_04110 | G2T0V0 | *Rh*Lnb136$_{I}$ | Reverse | GGTGGTGGTGCTCGAGGCCCGCCCGGTTTTCTGA |
| RHOM_04120 | G2T0V2 | *Rh*GLnbp112 | Forward | AGGAGATATACCATGGATGACTTTAAAAGAGGGACGTG |
| RHOM_04120 | G2T0V2 | *Rh*GLnbp112 | Reverse | GGTGGTGGTGCTCGAGGCCAATGTTATACCATTTAATCTCG |
| RHOM_04095 (AA36-454) | G2T0U7 | *Rh*LNBBP | Forward | TTTCAGGGCGCCATGGGTGCAGCTGAAACCAGCC |
| RHOM_04095 (AA36-454) | G2T0U7 | *Rh*LNBBP | Reverse | GACGGAGCTCGAATTCTTATTCACTAATGTTAAATTCAAC |
| ROSEINA2194_01898 (AA26-861) | C0FT31 | *Ri*Le$^{a/b}_{II}$136 | Forward | TTTCAGGGCGCCATGGGTAATGCAGGGACAACCT |
| ROSEINA2194_01898 (AA26-861) | C0FT31 | *Ri*Le$^{a/b}_{II}$136 | Reverse | GACGGAGCTCGAATTCTTATCTTCTGTAAAGCTCAAATTCT |
| ROSEINA2194_01899 (AA36-340) | C0FT32 | *Ri*Le$^{a/b}_{I}$136 | Forward | CAGCCATATGCTCGAGGGAGAAAATATTAAGATTTCCAAAG |
| ROSEINA2194_01899 (AA36-340) | C0FT32 | *Ri*Le$^{a/b}_{I}$136 | Reverse | CAGCCGGATCCTCGAGCTAATTCCATTTAATCGTATCG |
| ROSEINA2194_01885 | C0FT18 | *Ri*GLnbp112 | Forward | TTTCAGGGCGCCATGGGTAATAAAGAACACGGTGGAAGAGT |
| ROSEINA2194_01885 | C0FT18 | *Ri*GLnbp112 | Reverse | GACGGAGCTCGAATTCTTAAACAGCGTACCATTTAATCTCA |
| ROSEINA2194_01895 (AA23-470) | C0FT28 | *Ri*Le$^{a/b}$BP | Forward | TTTCAGGGCGCCATGGGAAATGCAAATACATCCGCAAACAC |
| ROSEINA2194_01895 (AA23-470) | C0FT28 | *Ri*Le$^{a/b}$BP | Reverse | GACGGAGCTCGAATTCTTATTGCGCAGTTTCTGAAACCTC |
| ROSEINA2194_01891/01890 | C0FT24/C0FT23 | *Ri*Fuc29 | Forward | TTTCAGGGCGCCATGGGGAGGACACCCGAAGAACAGA |
| ROSEINA2194_01891/01890 | C0FT24/C0FT23 | *Ri*Fuc29 | Reverse | GACGGAGCTCGAATTCTTATGATTCTTGATAAACCTCAA |
| ROSEINA2194_01889/01888/01887 | C0FT22/C0FT21/C0FT20 | *Ri*Fuc95 | Forward | TTTCAGGGCGCCATGGGGGATTTAAGTAAATATGATATTTG |
| ROSEINA2194_01889/01888/01887 | C0FT22/C0FT21/C0FT20 | *Ri*Fuc95 | Reverse | GACGGAGCTCGAATTCTTATCCTGTAATTTTTGCATTTC |
| ROSEINA2194_02198 (AA29-975) | C0FTX7 | *Ri*GH98 | Forward | TTTCAGGGCGCCATGGGCAAAACGGGATCAGAAT |
| ROSEINA2194_02198 (AA29-975) | C0FTX7 | *Ri*GH98 | Reverse | GACGGAGCTCGAATTCTTAAACTATATCAAAATACACAT |
| HMPREF0373_02965 | U2PDT9 | *Er*Lnb136 | Forward | TTTCAGGGCGCCATGGGAAAATTGTGTGAAAATCAGCAGG |
| HMPREF0373_02965 | U2PDT9 | *Er*Lnb136 | Reverse | GACGGAGCTCGAATTCTTAAATCAGATGGATTTCATTCTCC |
| HMPREF0373_02965 | U2PDT9 | *Er*Lnb136_Y145A | Forward | CGAAAACAGATCACCATGAGCCCTGGTAAACAGATCCTGG |
| HMPREF0373_02965 | U2PDT9 | *Er*Lnb136_Y145A | Reverse | CCAGGATCTGTTTACCAGGGCTCATGGTGATCTGTTTTCG |

[a] UniProtKB accession number

### Supplementary Figures:

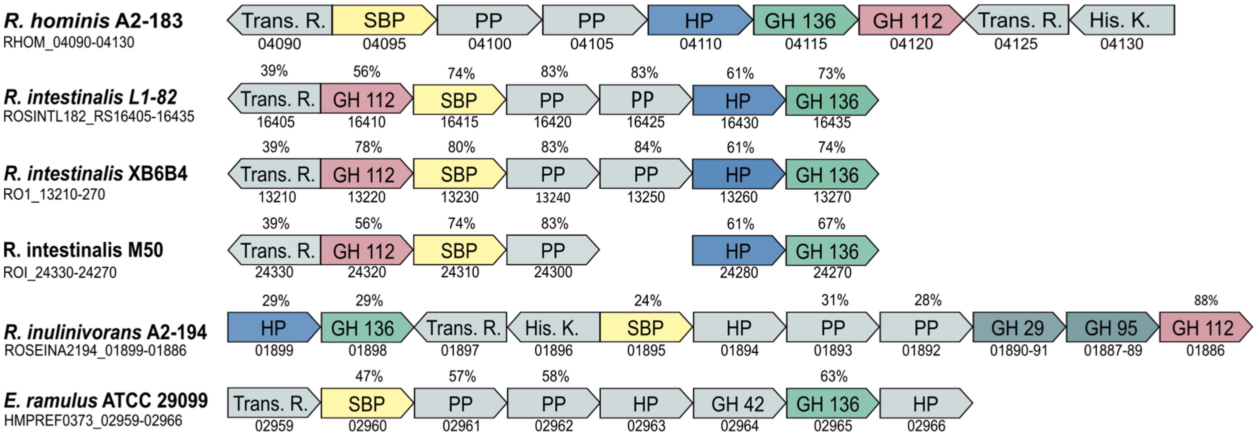


**Supplementary Fig. 1: The conservation of the core HMO utilization loci within *Roseburia* spp. and *Eubacterium ramulus*.** Gene locus IDs are below the genes, which are denoted according to their protein products: transcriptional regulator (Trans. R.); ABC transporter solute binding protein (SBP); ABC transporter permease protein (PP), hypothetical proteins (HP), glycoside hydrolase (GH) and histidine kinase sensory protein (His. K.). Sequence identities to the *R. hominis* A2-183 corresponding homologs are above the genes. Genes coding for GH136 family members were identified via the dbCAN database.

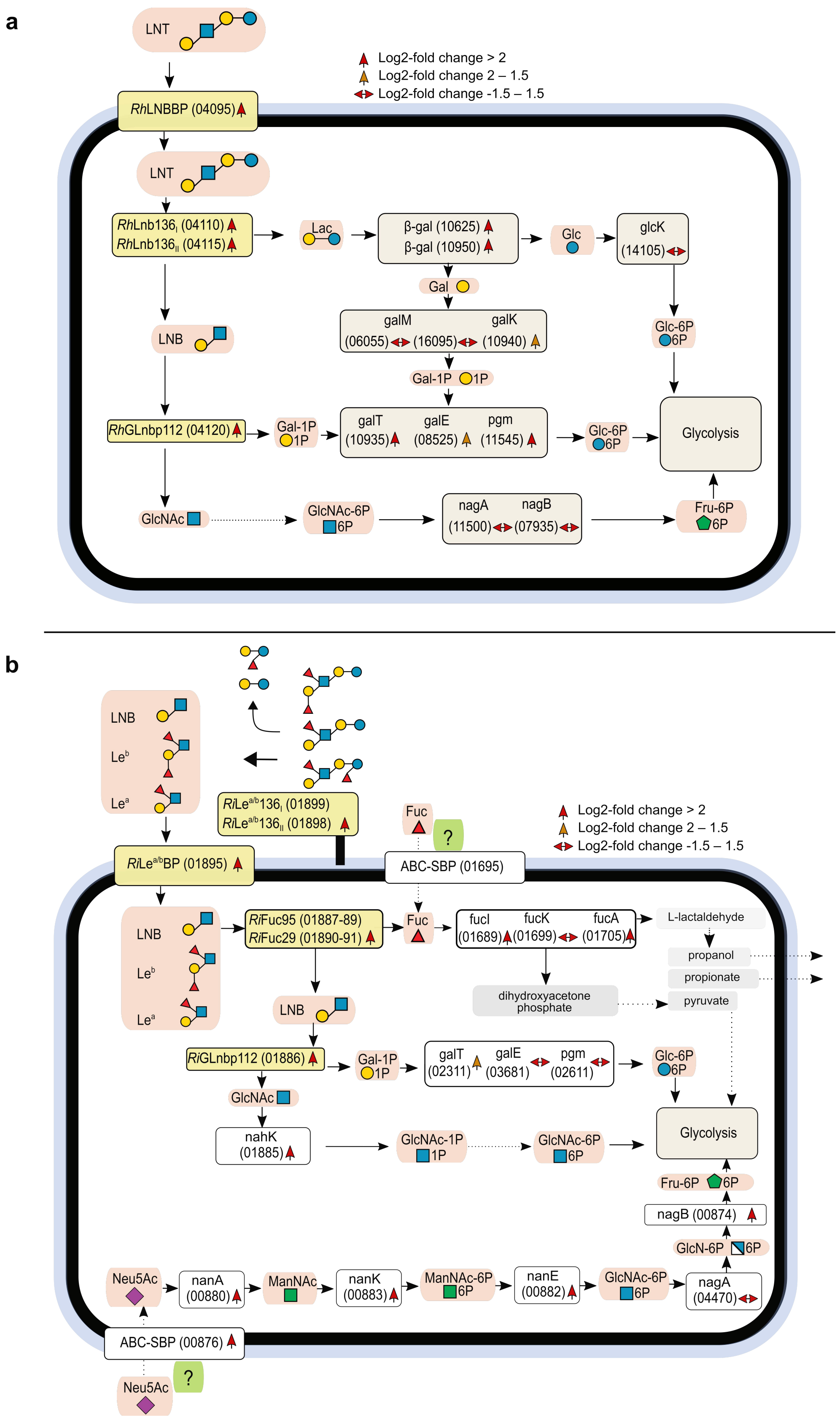


**Supplementary Fig. 2: Proposed HMOs core degradation pathways of *R. hominis* and *R. inulinivorans.* (a)** Proposed model for LNT utilization in *R. hominis* and **(b)** for fucosylated HMOs utilization in *R. inulinivorans,* based on proteomics data of cells grown on LNT (**a**, *R. hominis*) and HMOs from mother milk (**b**, *R. inulinivorans*) relative to glucose. Enzymatic steps suggested from literature and detected in proteomics data are indicated in solid arrows, steps suggested by literature but not detected in proteomics data are shown as dotted arrows. Characterized proteins in the present study are highlighted in yellow squares and Locus IDs of *R. hominis* (Rhom_xxxxx) and *R. inulinivorans* (Roseina2194_xxxxx) are abbreviated with the last numbers after the hyphen.

**a** *R. inulinivorans* L-fucose acid utilization locus

| Locus ID | Log2 fold change | | Protein | Annotation |
|---|---|---|---|---|
| | HMOs/Glc | Mucin/Glc | | |
| 01688 | | | | Transcriptional regulator |
| 01689 | 6.64 | 6.64 | fucI | L-fucose isomearse |
| 01690 | | | | Hypothetical protein |
| 01691 | 2.52 | | ABC-PP | ABC transporter, permease protein |
| 01692 | 3.04 | 6.64 | ABC-PP | ABC transporter, permease protein |
| 01693 | 5.18 | 4.72 | ABC-PP | ABC transporter, permease protein |
| 01694 | | | ABC-PP | ABC transporter, permease protein |
| 01695 | 5.59 | 6.64 | ABC-SBP | ABC, solute binding protein |
| 01696 | 1.65 | 2.1 | | FucU transport protein |
| 01697 | 0.37 | | | Hypothetical protein |
| 01698 | | | | Carbohydrate kinase |
| 01699 | 1.15 | 1.92 | fucK | L-fuculokinase |
| 01700 | 1.87 | | | Nucleotide binding domain ParA family protein |
| 01701 | 1.74 | 1.71 | | Hypothetical protein |
| 01702 | 1.46 | | | Hypothetical protein |
| 01703 | 2.58 | 5.2 | | BMC domain protein |
| 01704 | | | | Ethanolamine/Propanediol utilization protein |
| 01705 | 3.82 | 3.64 | fucA | L-fuculophasphate aldolase |
| 01706 | 2.43 | | | Hypothetical protein |
| 01707 | 3.97 | 3.35 | | Hypothetical protein |
| 01708 | 3.15 | 3.68 | | CoA dependent aldehyd dehydrogenase |
| 01709 | 3.83 | 4.12 | | Alcohol dehydrogenase |
| 01710 | 2.77 | 4.24 | | BMC domain protein |
| 01711 | 3.21 | 3.12 | | BMC domain protein |
| 01712 | 2.49 | 3.01 | | BMC domain protein |
| 01713 | 2.73 | 2.67 | | BMC domain protein |
| 01714 | 3.39 | 3.64 | | Phosphate propanoyltransferase |
| 01715 | 4.27 | | | Ethanolamine/Propanediol utilization protein |
| 01716 | 3.07 | 2.29 | | NADH dehydrogenase like subunit protein |
| 01717 | 2.11 | 2.52 | | BMC domain protein |
| 01718 | 2.65 | | | Transcriptional regulator |
| 01719 | 3.14 | 2.82 | | Propandiol dehydratase |
| 01720 | 2.64 | | | Pyruvate lyase |

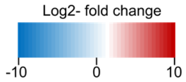


*R. inulinivorans N*-acetylneuraminic acid utilization locus

| Locus ID | Log2 fold change | | Protein | Annotation |
|---|---|---|---|---|
| | HMOs/Glc | Mucin/Glc | | |
| 00873 | | | | Transcriptional regulator |
| 00874 | 2.11 | 1.32 | nagB | glucosamine-6-phosphate deaminase |
| 00875 | 0.28 | 1.65 | | Transcriptional regulator |
| 00876 | 2.96 | 4.71 | | ABC transporter, solute binding protein |
| 00877 | | | | Hypothetical protein |
| 00878 | | | ABC-PP | ABC transporter, permease protein |
| 00879 | | | ABC-PP | ABC transporter, permease protein |
| 00880 | 2.57 | 5.02 | nanA | N-acetylneuraminate lyase |
| 00881 | 1.95 | 3.67 | YhcH | YhcH family protein |
| 00882 | 2.71 | 4.25 | nanE | N-acetylmannosamine-6-phosphate epimerase |
| 00883 | 2.89 | 4.65 | nanK | N-acetylmannosamine kinase |
| 00884 | 1.52 | 3.21 | AE | Acetylesterase |

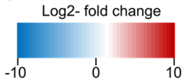


**Supplementary Fig. 3: *R. inulinivorans* upregulated L-fucose and *N*-acetylneuraminic acid utilization loci. (a)** Upregulation of L-fucose utilization cluster of *R. inulinivorans.* **(b)** Upregulation of putative *N*-acetylneuraminic acid utilization cluster of *R. inulinivorans* cells grown on purified HMOs from mother milk and mucin, respectively, relative to glucose (Glc). **(a,b)** The heatmaps depict Log2-fold changes of proteins expressed by cells grown on HMOs or mucin, respectively, relative to glucose. Locus numbers Roseina2194_xxxxx are abbreviated with the last numbers after the hyphen.

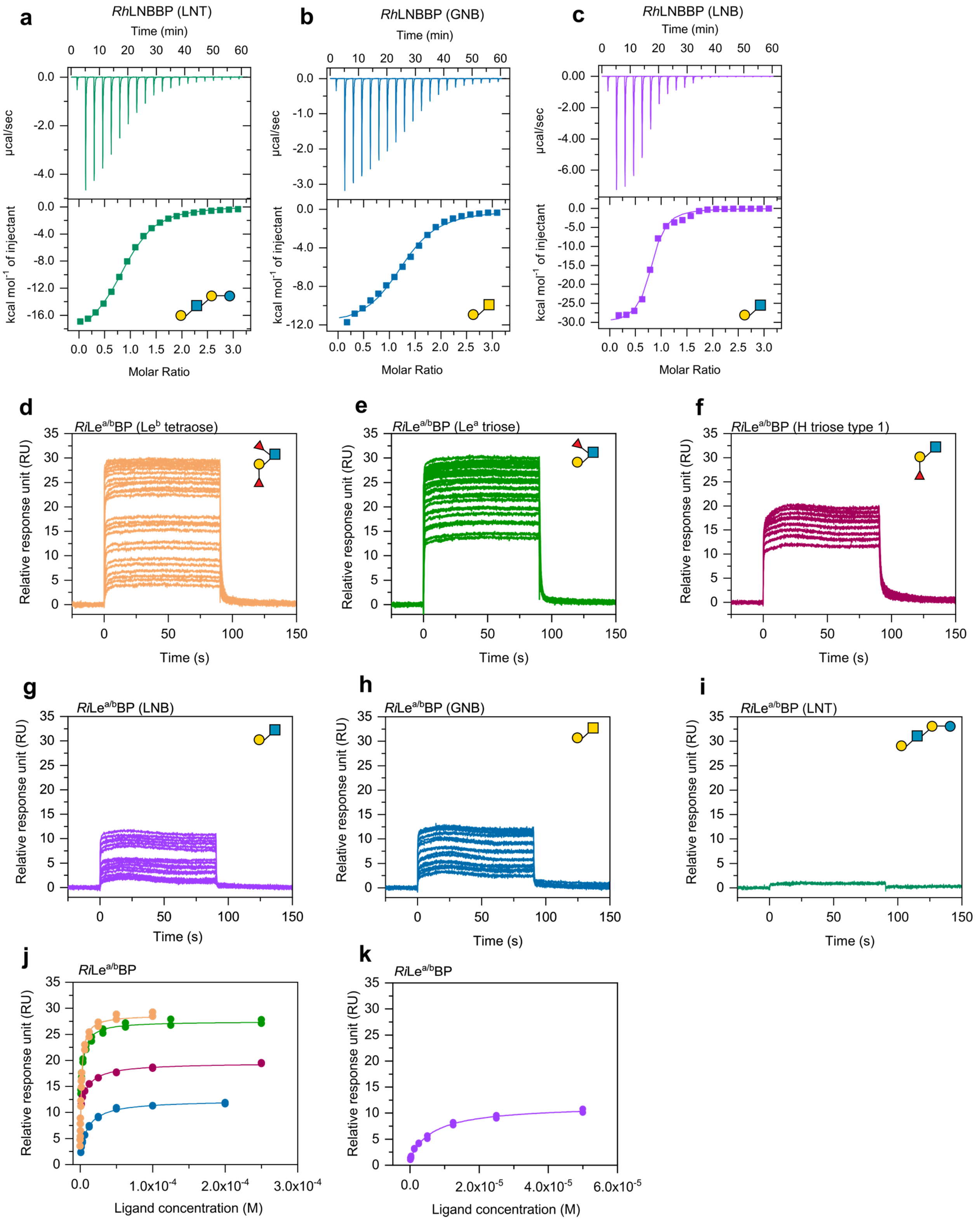


**Supplementary Fig. 4: Binding of *Rh*LNBBP and *Ri*Le$^{a/b}$BP to HMOs. (a-c)** ITC analysis of *Rh*LNBBP to selected HMOs. **(d-i)** Reference and blank corrected sensograms illustrating binding of selected HMOs to *Ri*Le$^{a/b}$BP. **(j-k)** One binding model fitted to the binding isotherms from the sensograms in (**d-i**). ITC and SPR experiments were performed as duplicates.

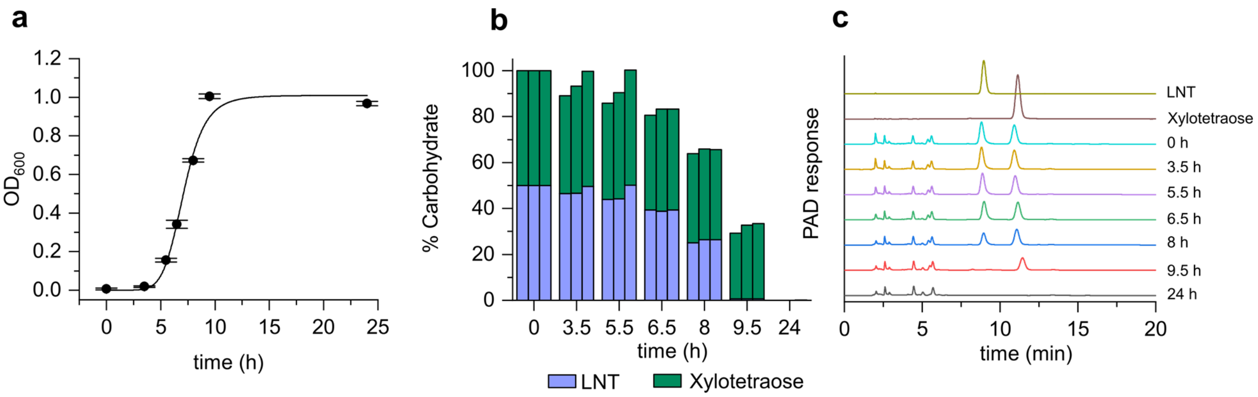


**Supplementary Fig. 5: Oligosaccharide uptake preference of *R. hominis* during growth on an equimolar LNT and xylotetraose mixture. (a),** Growth curve of *R. hominis* on YCFA supplemented with 0.5 % (w/v) of an equal mixture of xylotetraose and LNT. **(b),** Time course of relative percentages of xylotetraose and LNT in culture supernatants from (a) calculated based on HPAEC-PAD analyses as exemplarily represented in (c). **(c),** HPAEC-PAD chromatograms showing time course analysis of culture supernatants of *R. hominis* grown on YCFA supplemented with 0.5 % (w/v) of an equal mixture of xylotetraose and LNT. Observed peaks between 0 and 6 minutes are medium components. Growth experiment (a) and HPAEC-PAD analysis (b,c) were performed in triplicates.

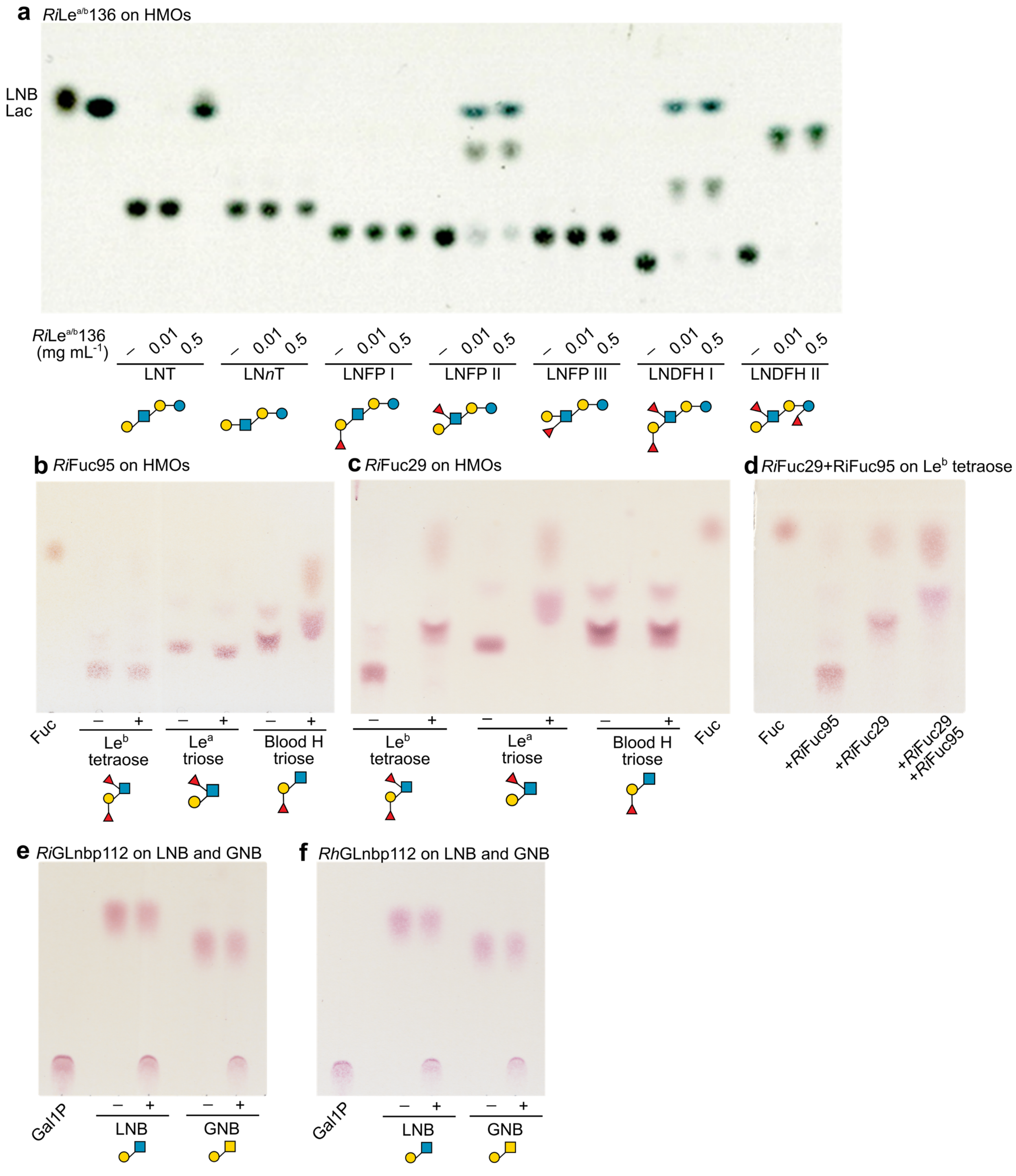


**Supplementary Fig. 6: Substrate preference of *Ri*Le$^{a/b}$136 and intracellular decomposition of GH136 degradation products in *Roseburia*.** **(a)**, Substrate preference of *Ri*Le$^{a/b}$136 towards HMOs; reactions with 0.01 or 0.5 mg mL$^{-1}$ of *Ri*Le$^{a/b}$136, respectively. **(b)**, Fucosidase activity of *Ri*Fuc95 on HMOs. **(c)**, Fucosidase activity of *Ri*Fuc29 on HMOs. **(d)**, Complete defucosylation of Le$^{b}$ tetraose by orchestral action of *Ri*Fuc29 and *Ri*Fuc95. Data show hindrance of *Ri*Fuc95 by α-(1→2)-linked L-fucose on Le$^{b}$ tetraose. **(e)**, Phosphorylase activity of *Ri*GLnbp112. **(f)**, Phosphorylase activity of *Rh*GLnbp112. **(a-f)**, +: reactions with enzyme, -: controls without enzyme. Analyses were performed in duplicates.

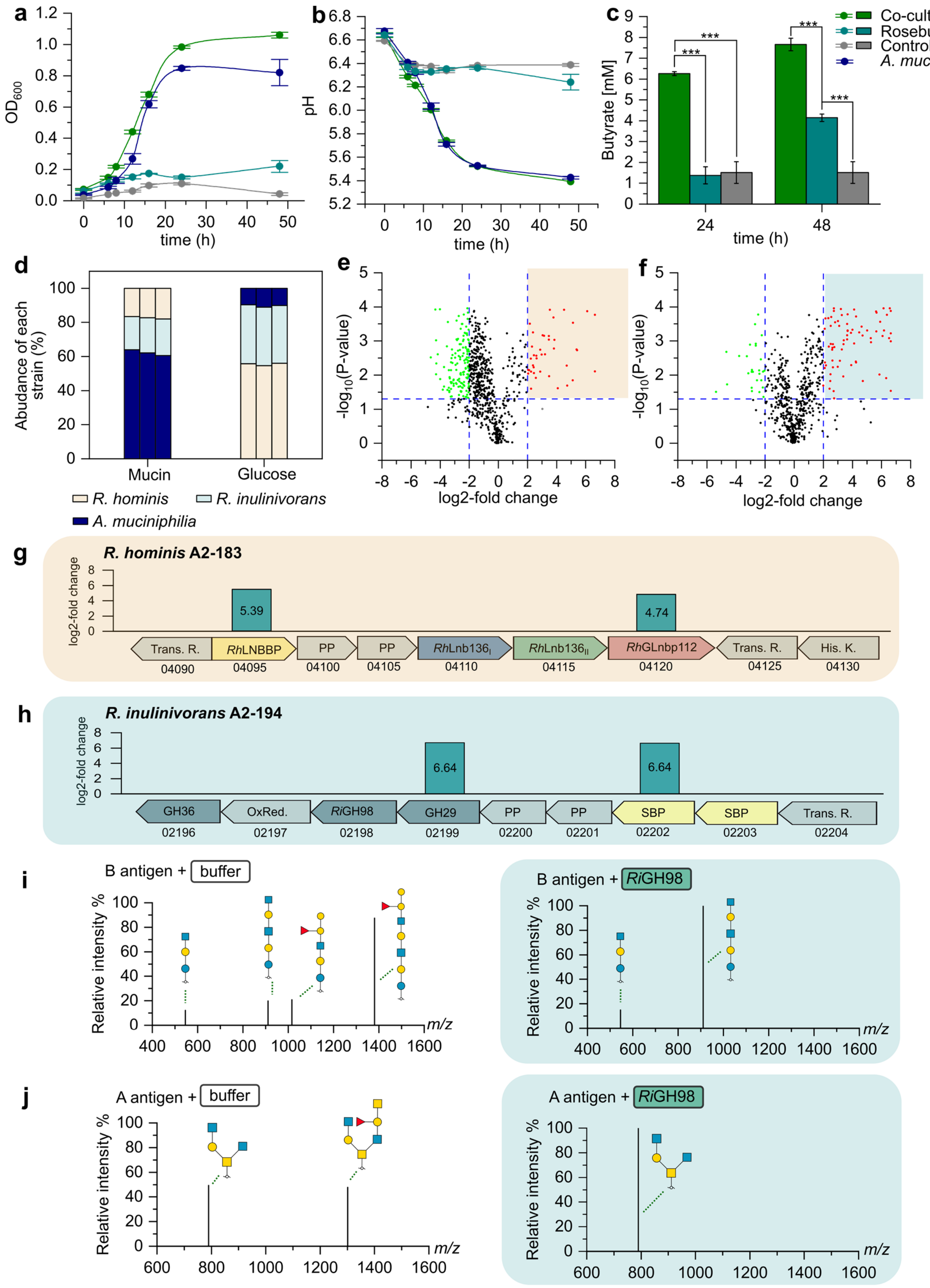


**Supplementary Fig. 7: Crossfeeding of *Roseburia* in *A. muciniphilia* co-cultures on mucin. (a-b)**, Growth of monocultures and co-cultures of *Roseburia* spp. and *A. muciniphila* on mucin. **(c)**, Butyrate in culture supernatants of monocultures and co-cultures as in (a) at 24h and 48h. **(d)**, Relative strain abundance during growth of co-cultures on mucin and glucose at 16 h determined based on MS/MS analyses. **(e)**, Volcano plot depicting upregulation pattern of proteins in *R. hominis* cells or **(f)**, in *R. inulinivorans* cells grown in mucin co-culture relative to glucose. **(g)**, Upregulated proteins in the core HMOs locus of *R. hominis* cells as in (e). Upregulated proteins in putative blood group utilization locus of *R. inulinivorans* cells as in (f). **(i-j)** Degradation of Blood group antigen A and B by *Ri*GH98 analyzed by nanoLC-MS. **(a-j)**, Growth cultures were performed in four replicates, proteomics analyses originate from biological triplicates and nanoLC-MS analyses were performed in duplicates. **(c)** Three asterisk (***) indicate a statistically significant difference at a level of $p < 0.001$

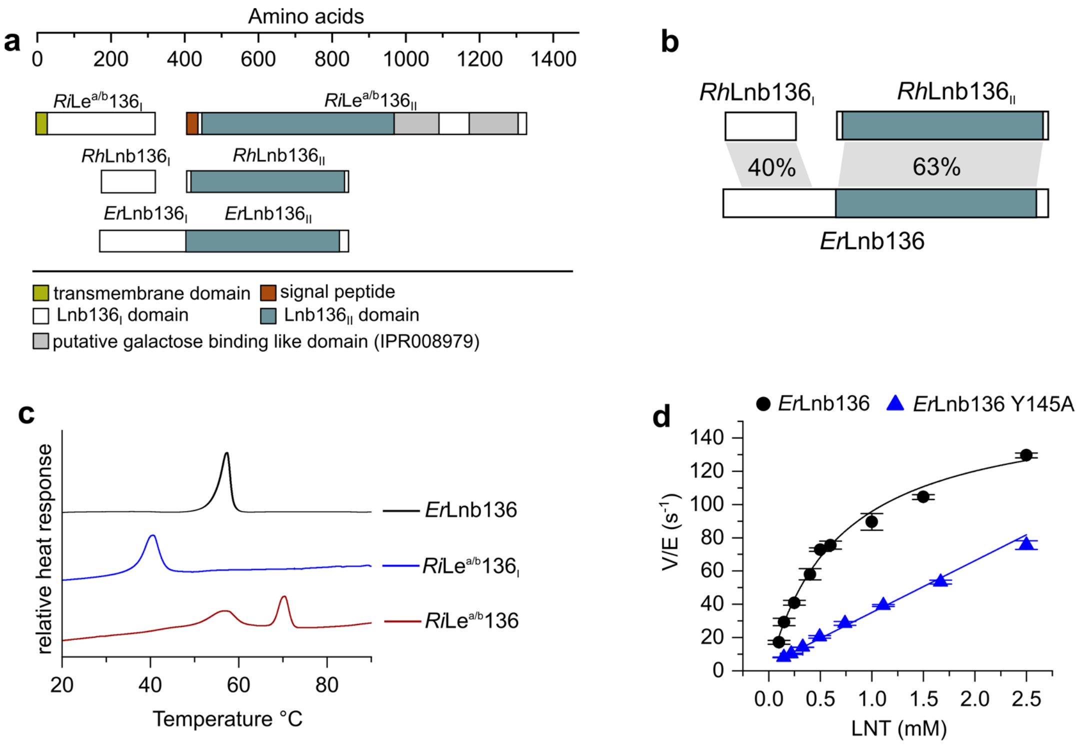


**Supplementary Fig. 8: Organization, stability and functional interactions of GH136 domains. (a)** domain organization of GH136 enzymes in *R. inulinivorans*, *R. hominis* and *E. ramulus*. **(b)** Amino acid sequence identities between the two GH136 domains *Rh*Lnb136$_{I}$ and *Rh*Lnb136$_{II}$ from *R. hominis* and *Er*Lnb136 from *E. ramulus*. **(c)** Differential scanning calorimetry thermograms showing the unfolding of *Er*Lnb136 and of *Ri*Le$^{a/b}$136$_{I}$ and *Ri*Le$^{a/b}$136. The unfolding of the two domains of ErLNb136 appears to overlap giving rise to a single asymmetric thermal transition consistent with the cooperative unfolding of the domains. By contrast, the unfolding of *Ri*Le$^{a/b}$136 features two well resolved transitions, the first is likely attributed to the unfolding of the *Ri*Le$^{a/b}$136$_{I}$ domain while the second is likely to be attributed to the unfolding of the remaining part of the protein including the *Ri*Le$^{a/b}$136$_{II}$ domain. **(d)**, Hydrolysis kinetics of *Er*Lnb136 and the mutant *Er*Lnb136 Y145A on LNT. DSC analyses (c) were performed as duplicates and kinetic measurements (d) were performed as triplicates.

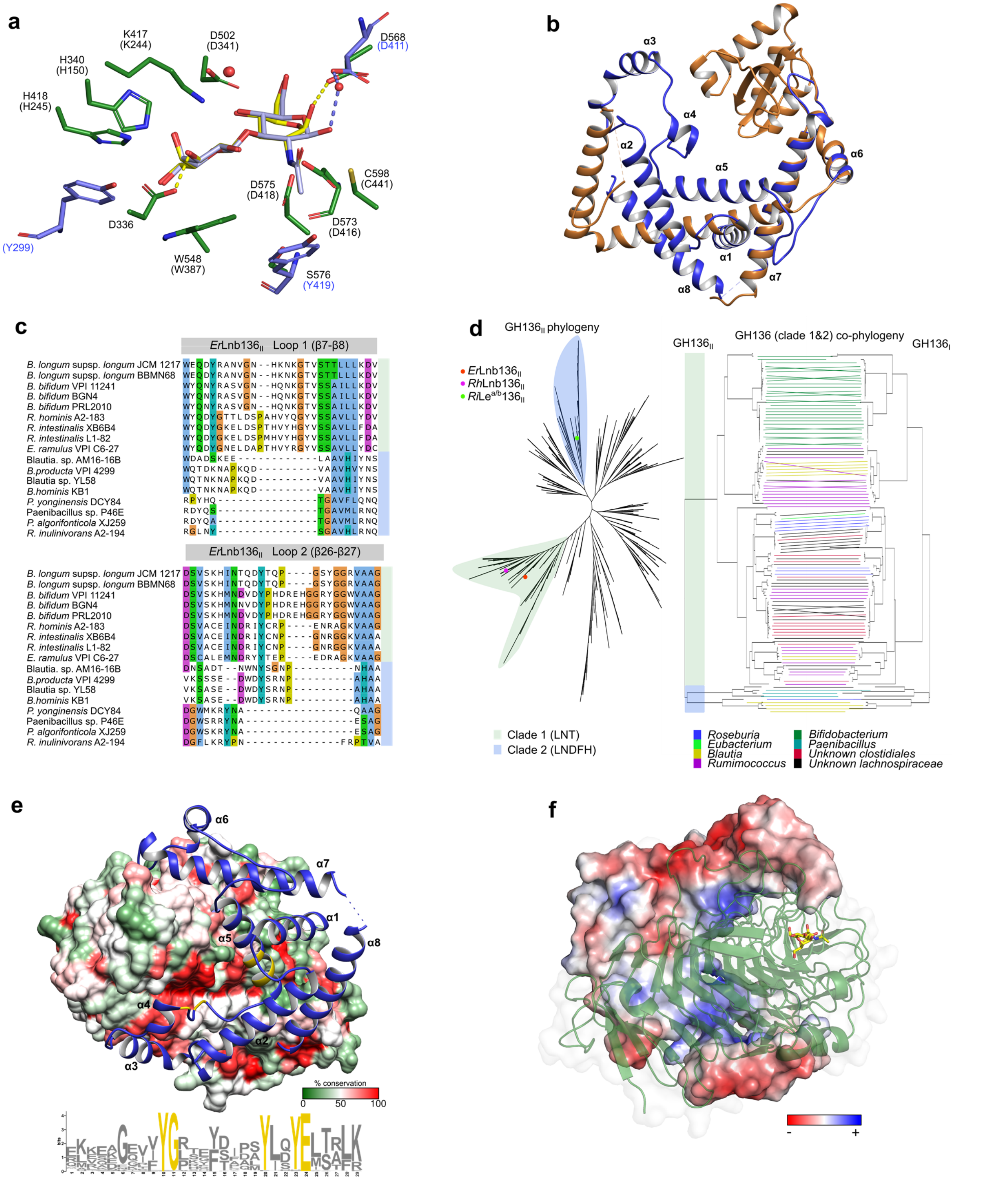


**Supplementary Fig. 9: Evolution of GH136 enzymes. (a)** Superimposition of LNB (yellow) bound in *Er*Lnb136 (green) and LNB (grey) in *Bl*LnbX from *B. longum* (light blue). Conserved residues are shown for *Er*Lnb136 and *Bl*LnbX (in parentheses). Residues Y299, Y419 and D411 of *Bl*LnbX that are variant compared to *Er*Lnb136 are shown in light blue to highlight differences in active site architecture and ligand binding. Water molecules are red spheres and hydrogen bonds are dashed lines in *Er*Lnb136 (yellow) and *Bl*LnbX (light blue). **(b)** Superimposition of *Er*Lnb136$_{I}$ (blue) and most related structural homolog 5EZ1 (chain A) from *Heliobacter pylori* (orange), highlighting the large differences in protein fold. **(c)** Partial amino acid sequence alignment of GH136$_{II}$ domains showing shortened loops around the active site in *R. inulinivorans* as compared to *Er*Lnb136 of *E. ramulus* **(d)** Phylogenetic tree of 985 GH136$_{II}$ sequences identified by BLASTP search of *Rh*Lnb136$_{II}$ or *Ri*Le$^{a/b}$136$_{II}$ against non-redundant database (sequences with an e-values < $10^{-10}$ are included). Tanglegram showing co-evolution of GH136$_{I}$ and GH136$_{II}$ domains across 117 selected sequences of GH136$_{II}$ phylogenetic tree clade 1 and clade 2. **(e)** Surface of *Er*Lnb136$_{II}$ colored by amino acid sequence conservation across 117 sequence as presented in (d) and cartoon presentation (blue) of *Er*Lnb136$_{I}$ with conserved residues (yellow) as identified from a sequence motif generated via the MEME suite from 117 GH136$_{I}$ sequences as in (d). **(f)** Electrostatic surface of *Er*Lnb136$_{I}$ and cartoon presentation of ErLnb136$_{II}$ (green).
